# Towards a Better Understanding of Batch Effects in Spatial Transcriptomics: Definition and Method Evaluation

**DOI:** 10.1101/2025.03.12.642755

**Authors:** Yingxin Zhang, Qingzhen Hou

## Abstract

**Background:** Spatial transcriptomics (ST) enables high-resolution mapping of gene expression within tissue slices, providing detailed insights into tissue architecture and cellular interactions. However, batch effects, arising from non-biological variations in sample collection, processing, sequencing platforms, or experimental protocols, can obscure biological signals, hinder data integration, and impact downstream analyses. Despite their critical impact, batch effects in ST datasets remain poorly defined and insufficiently explored. To address this gap, we propose a framework to categorize and define batch effects in ST and systematically evaluate the performance of ST methods with batch effect correction capabilities.

**Results:** We categorized batch effects in ST into four types based on their sources: (1) Inter-slice, (2) Inter-sample, (3) Cross-protocol/platform, and (4) Intra-slice. Seven ST integration methods—*DeepST, STAligner, GraphST, STitch3D, PRECAST, spatiAlign*, and *SPIRAL*—were evaluated on benchmark datasets from human and mouse tissues. Using metrics such as graph connectivity, kBET, ASW, and iLISI, we assessed both the preservation of biological neighborhoods and the effectiveness of these methods in batch correction. Additionally, we applied *STAligner* for downstream analysis to compare results before and after batch correction, further highlighting the importance of batch effect correction in ST analysis.

**Conclusion:** No single method is universally optimal. *GraphST, PRECAST, SPIRAL*, and *STAligner* performed well for same-platform integration, whereas *SPIRAL* and *STAligner* excelled in cross-platform settings. These findings highlight the need for robust and generalizable ST approaches with effective batch correction capabilities to facilitate the integration of multi-platform ST datasets in future research.

## 1 Introduction

Spatial transcriptomics (ST) provides high-resolution spatial mapping of gene expression across tissue slices, offering crucial insights into tissue architecture in both physiology and disease, as well as the cellular microenvironments that shape these structures[1][2]. As ST technologies continue to evolve, the scope of spatial transcriptomics data and analysis methods has expanded significantly[3][4]. New approaches have been developed to address the inherent challenges in spatially resolved gene expression profiling, significantly improving our understanding of complex biological systems[5][6]. These advances led to ST being recognized as the “Method of the Year” due to its transformative potential in understanding biological systems and diseases[7].

However, spatial transcriptomics analyses are frequently impacted by batch effects, which arise from differences in samples, technologies, measurement environments, or operational errors[8]. Batch effects obscure biological signals, reduce statistical power, and even result in misleading or biased conclusions, complicating data interpretation and integration. Addressing batch effects is therefore essential for reliable data integration and accurate biological interpretation, especially in large-scale, multi-platform, or multi-sample studies[9].

In ST, batch effects can exacerbate spatial structural distortions between tissue slices, further compromising the reliability of downstream analyses[10]. To mitigate these effects, two key methods have been developed in ST analyses: alignment and integration[11]. Alignment methods aim to reduce spatial variability by aligning data from different tissue slices into a common coordinate system (CCS). This transformation allows data from different slices to be aligned in a consistent spatial framework, enabling meaningful comparisons across slices or biological scales[12]. By standardizing spatial information, alignment methods facilitate the consistent interpretation of gene expression patterns and their relationships to biological functions at the whole-tissue level[8][13][14]. On the other hand, integration methods combine data from multiple sources into a unified feature space to provide a unified perspective, yielding more comprehensive, systems-level insights in biology. Integration enhances clustering accuracy and spatial domain identification, facilitating the study of tissue structure and function[8][15][16][17]. While alignment and integration are often performed together, batch effect correction primarily occurs during the integration process.

Spatial transcriptomics methods for alignment and integration can be categorized into three groups based on their batch effect correction capabilities and algorithms used:

### 1. Methods with inherent batch effect correction include

These methods directly incorporate batch effect correction into their spatial alignment and integration processes:

### Graph-based Methods

These methods align spatial coordinates and reduce batch variability using graph algorithms: *SLAT*[18] integrates graph adversarial matching and cross-modal embedding to align inter-batch spatial and transcriptional patterns. *STAligner*[19] employs graph attention autoen-coders and spatial neighbor graphs, leveraging attention-weighted embeddings to reduce batch noise. *GraphST*[20] combines PASTE[21] alignment with contrastive learning on shared neighborhood graphs for batch-invariant feature smoothing.

### 3D Integration Methods

*STitch3D*[22] adjusts gene expression by accounting for slice- and gene-specific effects, thereby effectively eliminating batch effects and integrating spatial information across multiple ST slices.

### Probabilistic Models

These methods use probabilistic frameworks or deep learning models to align joint embeddings and reduce batch effects: *PRECAST*[23] models batch effects as latent variables via Bayesian inference, enabling probabilistic spatial clustering. *MEML*[24] integrates Gaussian processes with mixed-effects models, treating batch effects as random intercept covariance matrices. *iSC*.*MEB*[25] jointly optimizes batch states and spatial clustering via hidden Markov random fields (HMRF) and expectation-maximization (EM) algorithms.

### Adversarial Training and Domain Adaptation

These methods leverage domain adversarial neural networks to map data from different batches to a shared latent space, mitigating batch effects: *SpatiAlign*[26] aligns domain distributions through contrastive loss optimization with maximum mean discrepancy (MMD). *DeepST*[27] trains graph neural network (GNN) autoencoders adversarially with gradient reversal layers (GRL) to generate batch-invariant embeddings. *SPIRAL*[28] employs GraphSAGE with Wasserstein distance-based domain adaptation, aligning feature distributions across batches.

### Multi-module Toolkits

These toolkits combine various methods, such as graph convolutional networks and adversarial training, for robust batch effect correction: *SPACEL*[29] unifies graph convolutional networks and adversarial domain confusion through hybrid loss optimization.

### Explicit Spatial Modeling Methods

These methods explicitly model spatial distributions and cell type characteristics to reduce batch effects while preserving biological context: *spaDo*[30] employs Dirichlet process mixture models with location-dependent priors to enforce spatial continuity while modelling batch effects. *scBOL*[31] aligns semantic anchors across batches using prototype contrastive learning in embedding space.

### 2. Methods requiring additional batch effect correction tools

These methods partially address batch effects and require external tools (e.g., Harmony) for full correction. Examples include: *STAGATE*[32], *SEDR*[33].

### 3. Methods without inherent batch effect correction

These methods focus on spatial domain detection or clustering but rely entirely on external tools for addressing batch effects. Examples include: *BASS*[34], *BayesSpace*[35], *SpaGCN* [36].

Although there is an increasing number of spatial transcriptomics analysis tools available, the definitions of batch effects remains vague, and the field lacks systematic assessments of batch effect correction during spatial transcriptomic alignment and integration. While existing benchmarks[11][37][38][39] often focus on clustering, alignment, or general data integration, they typically do not assess batch effect correction directly. This gap limits the ability to effectively evaluate, compare, and select appropriate methods for mitigating batch effects in spatial transcriptomics. To address this, our study proposes a framework for defining batch effects in spatial transcriptomics. We define batch effects in spatial transcriptomics based on data sources, technical differences, and spatial structural heterogeneity. Batch effects were categorized into four types:

1. **Inter-slice: technical variability across tissue slices from the same biological sample;**
2. **Inter-sample: non-biological variability across slices from different biological samples;**
3. **cross-protocol/platform: systematic variability arising from differences in experimental protocols or technological platforms;**
4. **Intra-slice: local technical variability within a single tissue slice**.

Then, we systematically evaluated several methods with batch effect correction capabilities: *DeepST, STAligner, GraphST, STitch3D, PRECAST, spatiAlign*, and *SPIRAL*. These methods were tested across multiple datasets with varying batch definitions, including the Human Dorsolateral Prefrontal Cortex (DLPFC) dataset[40] for inter-slice and inter-sample evaluations, the Human Breast Cancer dataset for inter-slice analysis, and the Coronal Mouse Brain and Mouse Olfactory Bulb (OB) datasets for cross-protocol and cross-platform assessments, respectively. Performance was assessed using four widely recognized metrics[41]: graph connectivity (GC), k-nearest neighbor batch effects test (kBET), integration local inverse Simpson’s Index (iLISI), and batch Average Silhouette Width (ASW). These metrics ensured that integration improved batch mixing while preserving biologically meaningful structures. Additionally, we applied *STAligner* to integrate 12 slices of the DLPFC dataset, demonstrating the critical role of batch effect correction in spatial domain identification, trajectory analysis, and differentially expressed gene (DEG) analysis.

## 2 Results

We extended the classical definition of batch effects to better capture the unique characteristics of spatial transcriptomic data, as shown in Fig. 1A. To visualize the effects of batch correction, we compared raw (pre-correction) data with corrected outputs using Uniform Manifold Approximation and Projection (UMAP) and spatial projections. Performance was quantified using four key metrics (Fig. 1B): GC measured the preservation of biologically meaningful structures, while kBET, iLISI, and ASW evaluated batch mixing effectiveness. The workflow of our study and the benchmarking metrics are summarized in Fig. 1.

**Fig. 1:**
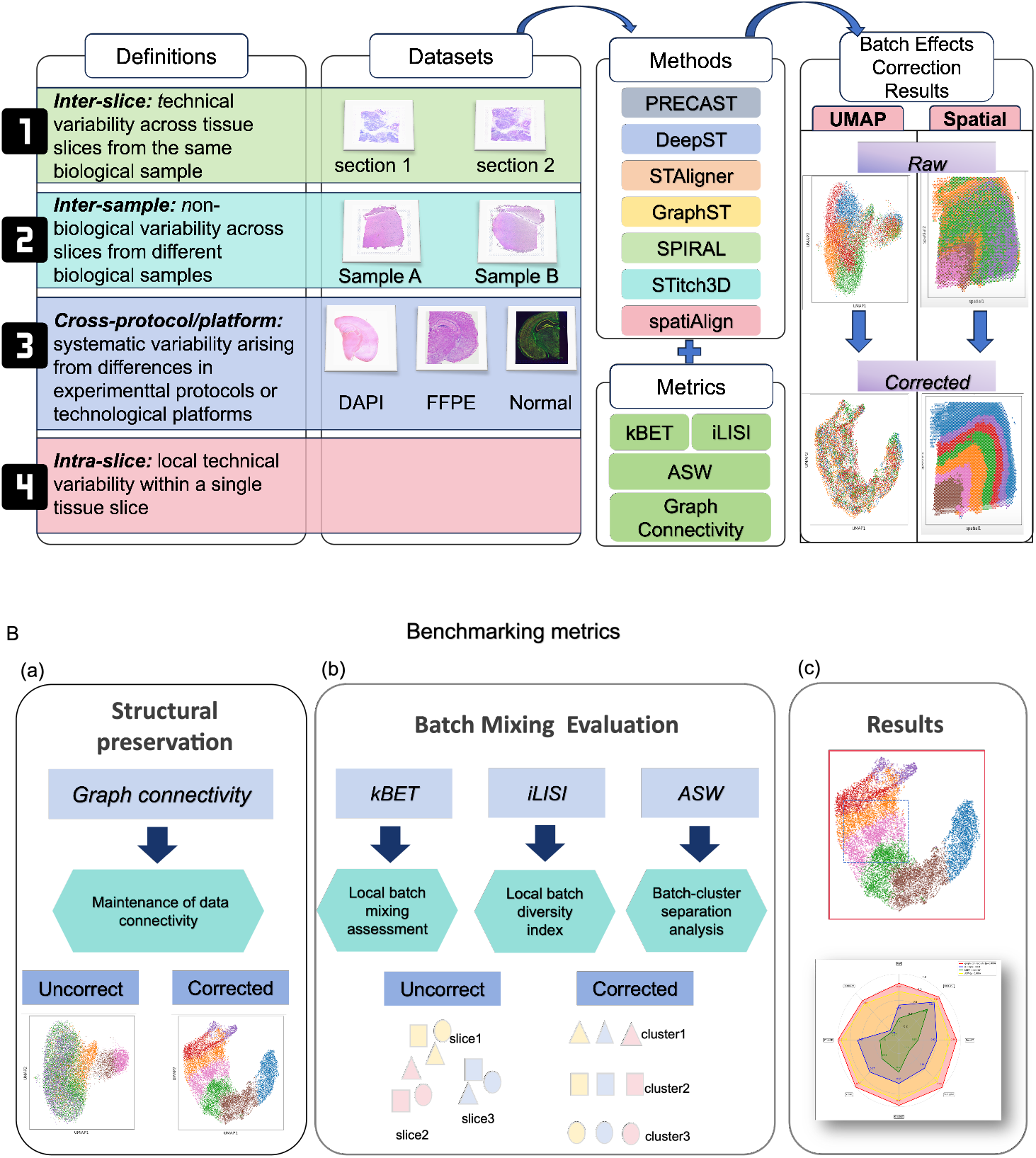
A: The workflow for evaluating batch effect correction of spatial transcriptomics in this study. This figure shows the framework of definitions, datasets, methods, and visualization of batch effect correction results. B: Descriptions of benchmarking metrics. (a) Graph connectivity (GC) assesses whether biologically meaningful groups (e.g., tissue layers) remain well-connected in integrated cross-batch graphs. It serves as an indirect measure of batch effects removal.(b) kBET quantifies how well batches (e.g., slices, samples) are mixed globally by comparing the local batch label distribution to the expected distribution under perfect mixing. iLISI evaluates diversity of batches in local neighborhoods; higher values indicate better fine-grained mixing of batches within small regions. The score ranges of kBET and iLISI is shown in the blue dashed box in the upper panel of Figure (c), corresponding to the regions where local batch mixing effects are evaluated. ASW measures the separation between batch labels and biological clusters, assessing global batch mixing. The score ranges of ASW is highlighted in the red box in the upper panel of Figure (c). The lower figure (c)shows the radar plot of the average values of different metrics in our study.

### 2.1 Definition of batch effects

In this study, we refined the classical definition of batch effects to better accommodate spatial context, considering additional sources of batch effects inherent to spatially resolved datasets.

#### 2.1.1 Inter-slice: technical variability across tissue slices from the same biological sample

Integrating consecutive or non-consecutive tissue slices from the same biological sample can introduce inter-slice batch effects, primarily driven by technical variability across experimental stages. Even slight differences in staining intensity, RNA quality, or sequencing depth between slices, despite originating from the same tissue, can introduce systematic biases. For instance, in Fig. 2A and 4A, our results showed that integrating non-consecutive slices “ABCD” or consecutive slices “AB” from the same DLPFC sample resulted in mismatched gene expression profiles and distorted spatial feature distributions. These biases would confound the interpretation of biologically relevant spatial patterns.

**Fig. 2:**
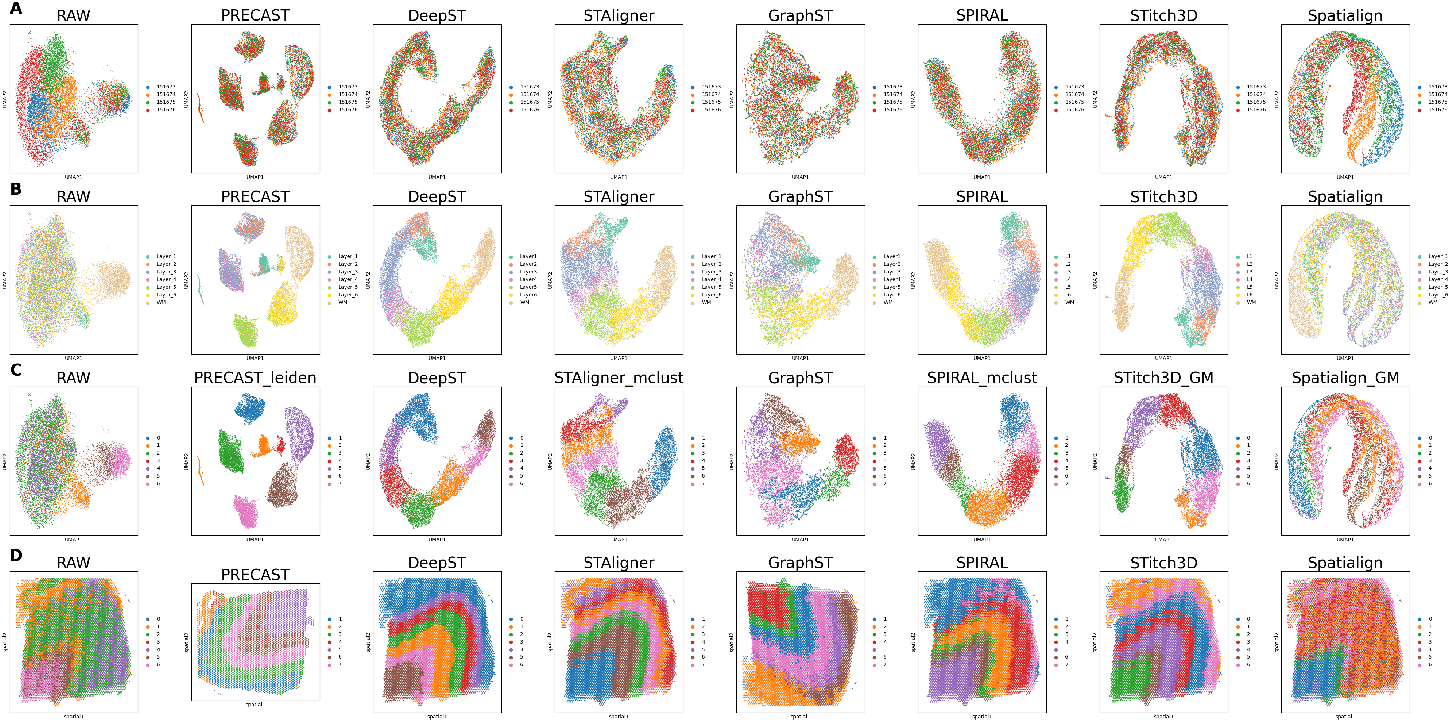
Visualization of the batch effect correction results from seven ST methods using DLPFC (Sample 1) dataset. The dataset contains four tissue slices: “sliceA: 151673”, “sliceB: 151674”, “sliceC: 151675”, and “sliceD: 151676”. Panels A-C show UMAP plots of uncorrected (RAW) data and seven ST methods (*PRECAST, DeepST, STAligner, GraphST, SPIRAL, STitch3D*, and *spatiAlign*). The UMAP plots are used to illustrate different perspectives. A: Colored by tissue slice indexes to demonstrate the mixture of different slices. B: Colored by manual annotations to assess whether tissue layers are clearly separated after correction. C: Colored by clustering results to show the separation of tissue layers after correction. Panel D demonstrates variations in spatial domain identification performance through clustering-label-derived metrics during non-consecutive slices integration.

#### 2.1.2 Inter-sample: non-biological variability across slices from different biological samples

Batch effects are more obvious when integrating tissue slices from multiple samples, particularly when those samples are processed across different experimental batches. Variations in reagents, equipment, operators, or processing time can introduce systematic biases that complicate data integration. For instance, uncorrected technical variability in the A region slices, integrated from DLPFC Samples 1–3, obscured biological distinctions in cortical layer markers, with batch effects surpassing true inter-layer variation (Fig. 6A).

#### 2.1.3. Cross-protocol/platform: systematic variability arising from differences in experimental protocols or technological platforms

Batch effects are also introduced when integrating data from different experimental protocols (e.g., FFPE vs. fresh frozen tissue) or when technological platforms differ (e.g., 10x Visium[42], Slide-seq[43], Stereo-seq[44]). Each platform has its own data format, sequencing depth, and capture efficiency, which introduces inconsistencies that complicate integration efforts. For example, slide-seq offers high spatial resolution but lower capture efficiency compared to platforms like Stereo-seq, which can result in inconsistent feature coverage across datasets. Similarly, FFPE tissue often suffers from RNA degradation compared to fresh frozen tissue, further complicating data integration and obscuring biological signals. In our observation, cross-protocol integration of coronal mouse brain (Fig. 8A) and cross-platform integration of mouse olfactory bulb (Fig. 10A) datasets introduced artificial regional expression biases, resulting in discordant anatomical alignment between different protocols or across distinct technological platforms.

#### 2.1.4 Intra-slice: local technical variability within a single tissue slice

Even within a single tissue slice, spatial heterogeneity can introduce batch-like effects. Variations in tissue location, experimental conditions, or technical discrepancies (e.g., uneven staining or fixation) can result in location-dependent systematic variations, often referred to as positional batch effects. These variations can lead to inaccurate spatial gene expression maps, distorting biological gradients or introducing clustering artifacts that obscure true spatial patterns[8]. Integrating data from different regions of the same slice requires careful consideration of these positional biases to ensure accurate representation of spatial relationships. However, further analysis to define and address these positional biases has not been conducted due to technical limitations.

### 2.2 Benchmarking

After defining the batch effects, we evaluated the performance of seven ST methods with batch effect correction capabilities: *DeepST, STAligner, GraphST, STitch3D, PRECAST, spatiAlign*, and *SPIRAL*, following the recommended preprocessing pipelines for each method. Specifically, we compared their ability to correct batch effects across three definitions: inter-slice, inter-sample, and cross-protocol/platform.

Initially, for each sample, the raw data, including gene expression matrices, spatial coordinates, and metadata were used for normalization, filtering to retain high-variance features, and annotation with batch and ground-truth labels (where available). Subsequently, these seven ST methods were applied to correct batch effects and generate embeddings. These embeddings were visualized by UMAP to evaluate two aspects: global batch mixing (determined by batch identifiers) and cluster separation (assessed against biological labels). Concurrently, spatial fidelity was validated by projecting domain clusters and corrected features onto the original tissue coordinates, ensuring that the resulting spatial patterns were biologically coherent. Finally, we computed kBET (local batch balance), iLISI (batch diversity), ASW (global mixing with biological coherence), and GC (biological neighborhood integrity) to holistically evaluate batch correction performance. In this evaluation, higher scores indicate better performance.

We used 10X Genomics Visium datasets of the DLPFC and human breast cancer (HBC) to evaluate the effectiveness of various methods in correcting batch effects during the integration of tissue slices from the same sample. The DLPFC dataset includes three neurotypical human samples (Samples 1–3). Each sample has two pairs of “spatial replicates” composed of 10 *µ*m adjacent slices, with the second pair located 300 *µ*m posterior to the first. These four slices, labeled A, B, C, and D, represent two replicates (AB and CD) with BC spaced 300 *µ*m apart. The slices were manually annotated in the original study into seven domains: six cortical layers (Layer_1_ to Layer_6_) and white matter (WM). DLPFC Sample 1 and DLPFC Sample 3 include all seven domains, while Sample 2 includes only five. Leveraging these manual annotations as ground truth, we assessed the performance of seven spatial transcriptomics methods with batch effect correction capabilities. The HBC dataset consists of two sections, Block A Section 1 and Section 2. Each section comprises 10 *µ*m thick cryosectioned slices of fresh frozen Invasive Ductal Carcinoma (IDC) breast tissue obtained from BioIVT Asterand, prepared according to the Visium Spatial protocol.

For Definition 1 (inter-slice), slices A, B, C, and D from each of the three DLPFC samples (Samples 1–3) were individually integrated to analyze non-consecutive slices. To further evaluate each method’s ability to correct batch effects, the HBC dataset (named as Sample 4) was also included in the analysis of non-consecutive slices. In contrast, slices A and B from Sample 1 were referred to as Sample 5, and used for the analysis of consecutive slices.

For Definition 2 (inter-sample), three slices from the A region of DLPFC Samples 1–3 were integrated to create a new dataset, DLPFC Sample 6, to validate the batch effect correction of different methods.

For Definition 3 (cross-protocol/platform), we used two datasets—coronal mouse brain (Sample 7) and mouse olfactory bulbs (Sample 8)—generated with varying experimental protocols and crossplatform ST data. First, coronal mouse brain datasets sequenced via 10X Visium technology under three experimental conditions were evaluated: FFPE tissue with H&E staining, fresh frozen tissue with H&E staining, and fresh frozen tissue with immunofluorescence (IF) staining. These datasets were labeled “10X_FFPE,” “10X_Normal,” and “10X_DAPI,” respectively. These protocols allowed us to assess the impact of tissue preparation and staining methods. Next, we conducted a comprehensive comparative analysis of ST data from mouse olfactory bulbs (OB) generated by three platforms: 10X Visium, Stereo-seq, and Slide-seq V2, with spatial resolutions of 50 *µm*, 35 *µm*, and 10 *µm*, respectively. These datasets were referred to as “10X,” “stereo-seq,” and “SlideV2” in this study. To ensure consistency, preprocessed data from the 10X platform was converted into AnnData objects using the SPIRAL method. These AnnData objects were then used for downstream integration analyses. Finally, we selected a total of 12 slices from DLPFC Samples 1–3 for downstream analysis using *STAligner*. An overview of the datasets is presented in Table 1.

**Table 1:** Suggestion of ST methods in different batch effect correction (Top1, Top2)

|  | Types | Datasets | Top1 | Top2 |
| --- | --- | --- | --- | --- |
| Inter-slice | Non-consecutive | DLPFC<br>(Sample 1,2,3) | *GraphST, STAligner* | *STitch3D* |
|  | Non-consecutive | HBC<br>(Sample 4) | *GraphST* | *PRECAST* |
|  | Consecutive | DLPFC<br>(Sample 5) | *GraphST* | *SPIRAL, STitch3D* |
| Inter-sample | Inter-sample | DLPFC<br>(Sample 6) | *PRECAST* | *STAligner, GraphST* |
| Cross-protocol<br>/platform | Cross-protocol | Coronal mouse brain<br>(Sample 7) | *STAligner* | *SPIRAL* |
|  | Cross-platform | Mouse OB<br>(Sample 8) | *SPIRAL* | *STAligner* |

This table summarizes the top two methods for each definition, based on UMAP visualization, batch effect assessment metrics, and datasets.

#### 2.2.1 Definition 1: technical variability across tissue slices from the same biological sample

For Definition 1, we tested two situations: non-consecutive and consecutive tissue slices from the same biological sample. For non-consecutive slices, Fig. 2 compared the raw data and batch effects-corrected results through UMAP visualization and spatial projections. Spatial transcriptomic spots, representing discrete tissue regions with spatially resolved gene expression profiles, were seperately colored by slice indexes, manual annotations, and clustering outcomes. Fig. 2D showed spatially coherent domains before and after batch effect correction. As shown in Fig. 2, the uncorrected data exhibited strong batch effects, characterized by systematic technical variations between slices from the same tissue. These batch effects distorted gene expression patterns, leading to mixed or misaligned features.

After applying the seven methods, most successfully integrated the slices and corrected batch effects, except for *spatiAlign*. The visualization of the annotated layers across these methods and raw data (Fig.2B) revealed several key findings. *STAligner, STitch3D, SPIRAL*, and *DeepST* successfully separated all seven layers and removed batch effects. *PRECAST* struggled to differentiate layer_2_, layer_4_ and layer_3_, failing to distinguish them well. Moreover, by comparing Fig.2 B and C, we found that the layer distributions obtained from clustering by *STAligner, GraphST*, and *SPIRAL* were generally consistent with the true layer distributions. In the spatial domain recognition plots (Fig. 2D), six methods clearly identified the spatial domains, while *spatiAlign* showed less distinct and more obscure layers.

As shown in Fig. 3, we evaluated batch effect correction capabilities using GC, iLISI, kBET, and ASW. Methods such as *STAligner, STitch3D, SPIRAL, GraphST* and *DeepST* showed high GC (*>*0.99), indicating effective batch effects removal and integration. The results were consistent with Fig. 2C, where spots from the same layer were clustered together. Among these methods, *GraphST* excelled in iLISI (0.83), highlighting its ability to maintain local similarity despite batch effects. *PRECAST* and *GraphST* also performed well in kBET, demonstrating optimal batch mixing. In contrast, *spatiAlign* faced some challenges in batch integration, as reflected in its lower GC and ASW scores, suggesting that batch effects were not fully minimized. This was also evident in the layer separation, where spots from the same layer showed less distinct separation (Fig. 2C).

**Fig. 3:**
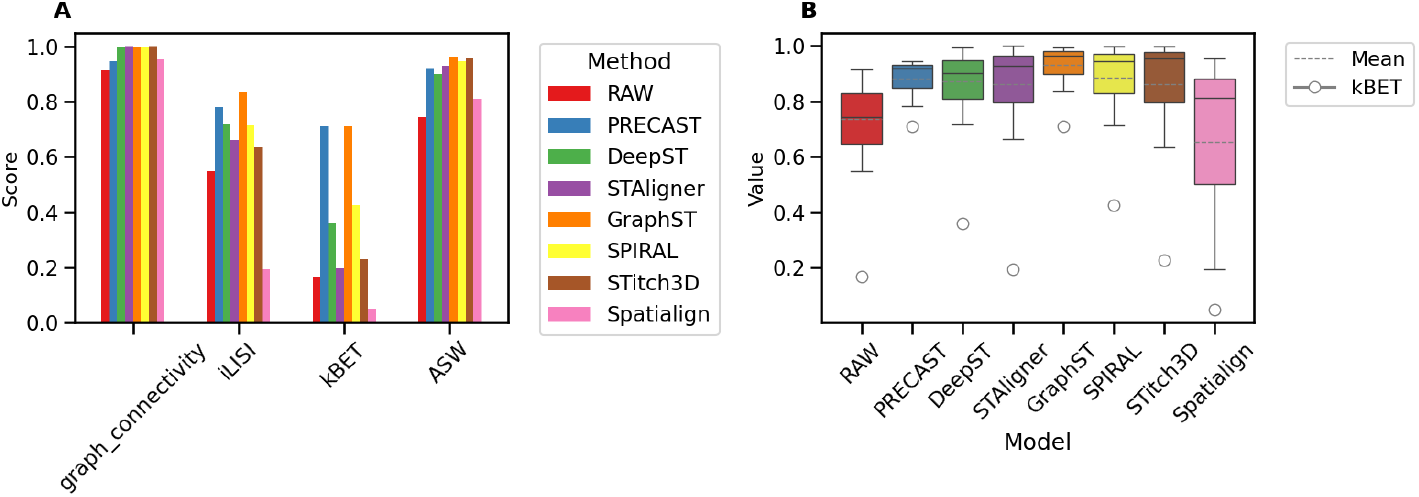
The figure compares seven methods using DLPFC (Sample 1) based on metrics related to batch effect correction in spatial transcriptomics. A presents the scores of each method on four metrics (GC, iLISI, kBET, ASW); B shows the mean values and quartile distributions of three metrics, (GC, iLISI, ASW), and kBET in circle.

Overall, *GraphST* achieved the best performance across all metrics for batch effects removal in non-consecutive slices, demonstrating excellent batch effects reduction while preserving biological structures. Methods such as *STAligner, STitch3D, SPIRAL*, and *DeepST* also performed well, with solid batch effect correction, as indicated by average scores above 0.70.

We also evaluated these methods for the integration of non-consecutive slices using Sample 2 and Sample 3 from the DLPFC dataset, as well as the HBC dataset, all under Definition 1. *GraphST, PRECAST, SPIRAL*, and *STAligner* consistently demonstrated effective layer separation (Supplementary File, Figures S1–S4), with identical cluster labels grouping together and batch effects being mitigated. These methods also preserved spatial domain structures well, highlighting their robustness in inter-slice spatial transcriptomics. In the HBC dataset (Supplementary File, Figures S5–S6), *GraphST* and *PRECAST* exhibited particularly effective batch effect correction, as reflected in their high iLISI and kBET scores. Notably, *GraphST* minimized batch effects while maintaining excellent spatial preservation, further underscoring its effectiveness across different datasets.

For consecutive slices, similar analyses were conducted (Fig. 4). The uncorrected data showed significant spatial overlap. Methods like *STAligner, GraphST, SPIRAL, DeepST, STitch3D*, and *PRECAST* successfully identified spatial domains by maintaining distinct layer separation. However, *spatiAlign* faced challenges in distinguishing multiple tissue layers, with the layers appearing more blended, leading to relatively lower performance compared to other methods. According to the metrics, *GraphST* and *SPIRAL*were the most successful in minimizing batch effects, facilitating better integration across datasets. *GraphST* performed particularly well across all metrics, including GC (0.99), ASW(0.94), iLISI(0.94), and achieving the highest average score (Fig. 5B), suggesting better integration of inter-slice data. *DeepST, PRECAST*, and *STitch3D* also demonstrated effective batch effects removal, which were consistently in Fig.4C, with average metric scores above 0.90. On the other hand, *spatiAlign* showed relatively weaker performance (Fig. 5).

**Fig. 4:**
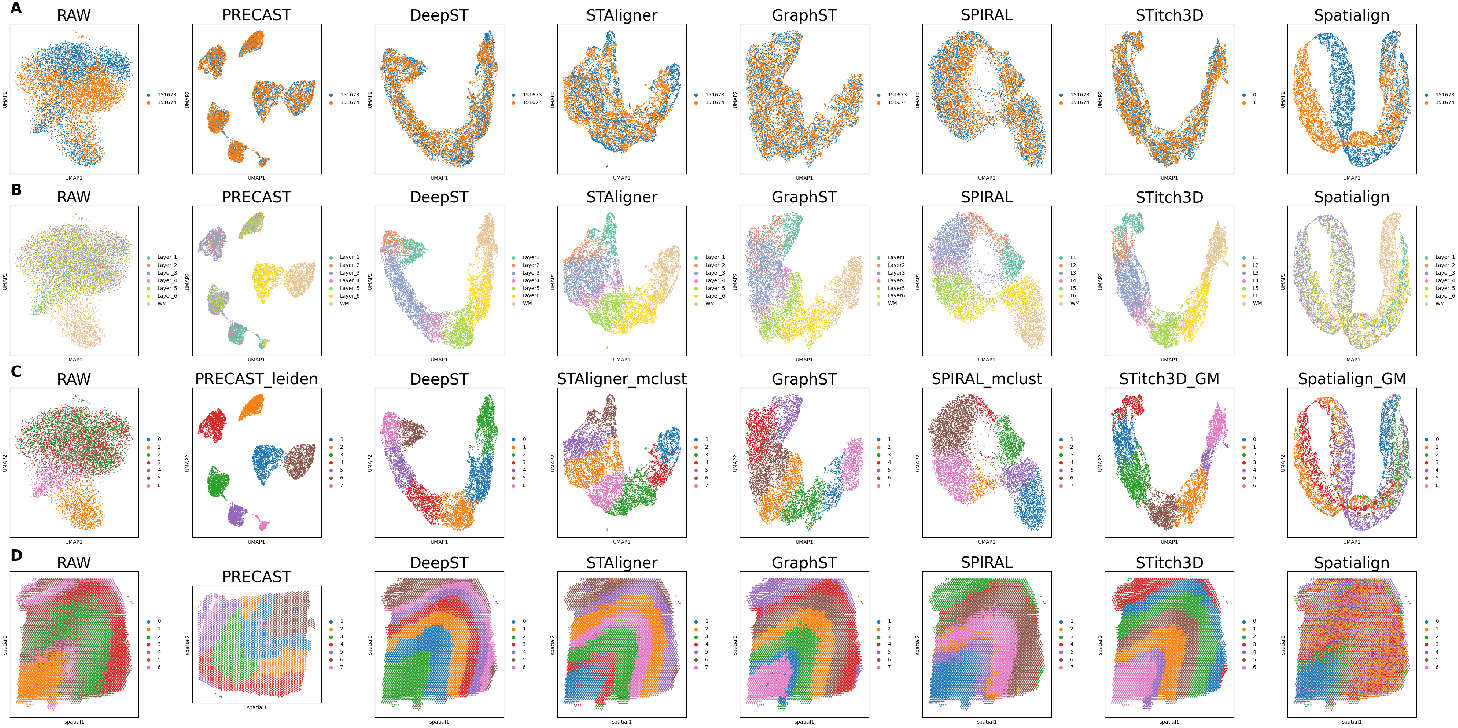
Visualization of the batch effect correction results from seven ST methods using DLPFC (Sample 5) dataset. The dataset includes two tissue slices: “sliceA: 151673”, “sliceB: 151674”. Panels A-C show UMAP plots of uncorrected (RAW) data and seven ST methods (*PRECAST, DeepST, STAligner, GraphST, SPIRAL, STitch3D*, and *spatiAlign*). Each UMAP plot is colored by three different setups, A: slice indexes, B: manual annotations, and C: clustering results. Panel D visually compares spatial domain identification performance during consecutive slices integration, employing clustering-label-derived metrics.

**Fig. 5:**
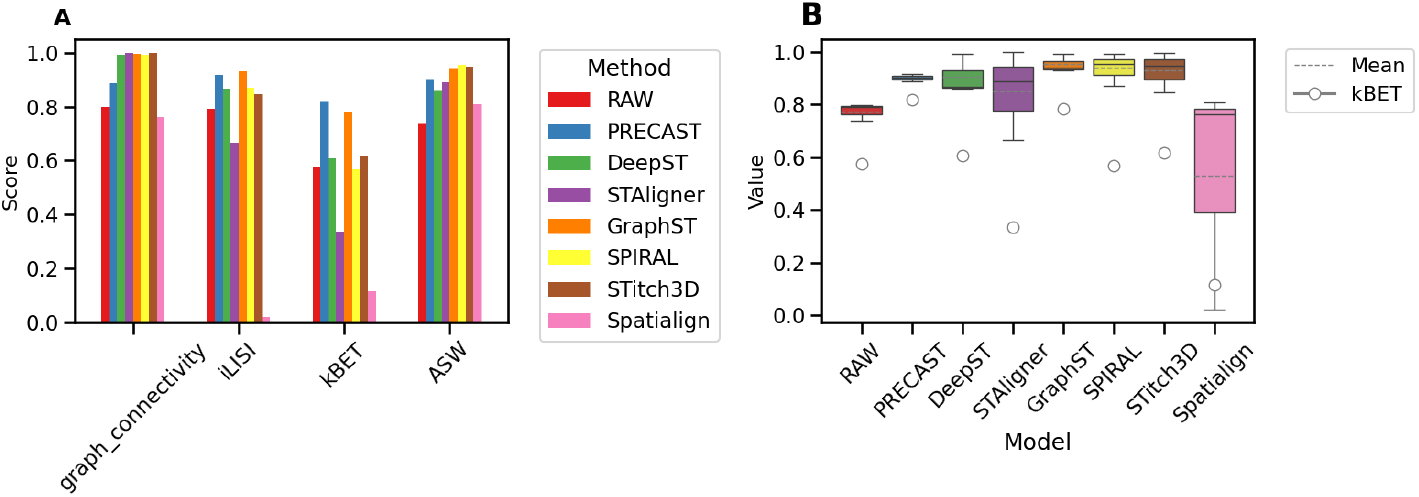
The figure compares seven methods using DLPFC (Sample 5) based on metrics related to batch effect correction in spatial transcriptomics. A presents the scores of each method on four metrics (GC, iLISI, kBET, ASW); B shows the mean values and quartile distributions of three metrics (GC, iLISI, and ASW), and kBET in circle.

In summary, *GraphST, STAligner, SPIRAL, DeepST, STitch3D*, and *PRECAST* demonstrated comparable performance in correcting batch effects associated with the integration of inter-slice data.*GraphST* exhibited consistent effectiveness in batch effect correction for both non-consecutive and consecutive tissue slices. Its strong performance in both batch correction and spatial preservation suggests it as a reliable option for addressing inter-slice batch effects.

#### 2.2.2 Definition 2: non-biological variability across slices from different biological samples

In Definition 2, we compared UMAP dimensionality reduction and spatial projection in the cross-sample slice analysis of DLPFC (Sample 6). We observed the methodological differences in mitigating batch effects and maintaining spatial structure (Fig. 6). Among the evaluated methods, *STAligner, DeepST, SPIRAL, PRECAST* and *STitch3D* successfully preserved the seven-layer hierarchical structure in the latent embeddings. These methods demonstrated strong capabilities in retaining spatial domain structures. Notably, *STAligner* exhibited superior spatial coherence in the UMAP visualizations, achieving tight cluster separation while retaining biologically meaningful spatial relationships. In contrast, *PRECAST* showed scattered representations of `layer`_4_ and `layer`_3_, suggesting that batch effects were only partially mitigated. Similarly, *GraphST* performed less effectively, with overlapping regions in `layer`_2_, `layer`_3_ and `layer`_4_. This overlap obscured layer boundaries and led to poor layer separation in the latent space.

**Fig. 6:**
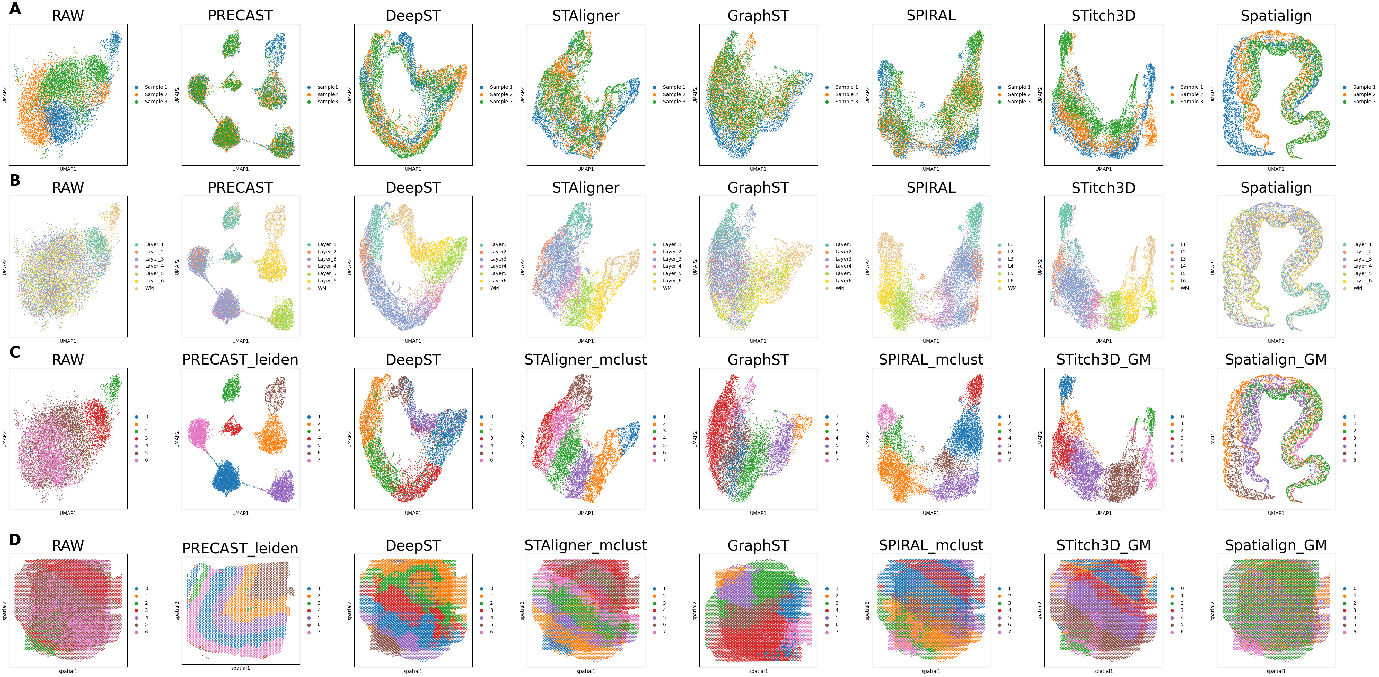
Visualization of the batch effect correction results from seven ST methods using DLPFC (Sample 6) dataset. The dataset includes three tissue slices from three samples: “Sample 1:151573”, “Sample 2: 151569”, and “Sample 3: 151507”. Panels A-C show UMAP plots of uncorrected (RAW) data and seven ST methods (*PRECAST, DeepST, STAligner, GraphST, SPIRAL, STitch3D*, and *spatiAlign*). Different colors are assigned based on A: sample indexes, B: manual annotations, and C: clustering results. Panel D compares spatial domain identification performance using clustering-label-derived visualizations during inter-sample integration.

Quantitative metrics such as GC, iLISI, kBET, and ASW (Fig. 7) were used to complement the visual results. In terms of overall performance, *PRECAST* achieved the highest mean score of 0.77, demonstrating its effectiveness in reducing batch effects. This was further supported by the distinct separation of tissue layers and improved batch mixing observed in Fig. 6A-C, suggesting effective batch effect correction. Both *GraphST* and *SPIRAL* also performed well, with mean scores of 0.72 and 0.71, respectively. Although *STAligner* had a relatively lower mean score, it showed more effective batch effect correction compared to *GraphST*, as shown in Fig. 6. However, when considering all factors, including GC and kBET, the overall performance of *GraphST* and *STAligner* appears to be quite similar, suggesting comparable capabilities in inter-sample integration.

**Fig. 7:**
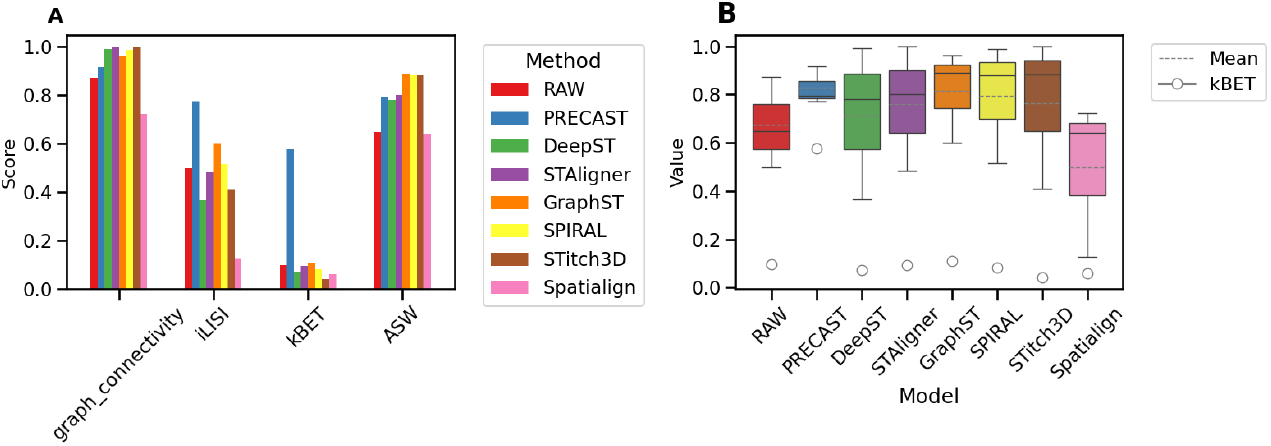
The figure compares seven methods using DLPFC (Sample 6) based on metrics related to batch effect correction in spatial transcriptomics. A presents the scores of each method on four metrics (GC, iLISI, kBET, ASW); B shows the mean values and quartile distributions of three metrics (GC, iLISI, and ASW), and kEBT in circle.

#### 2.2.3 Definition 3: systematic variability arising from differences in experimental protocols or technological platforms

##### Cross-protocol Integration

In the analysis of Definition 3, given that the inputs to *STitch3D* consist of multiple ST slices and a matched single-cell RNA sequencing (scRNA-seq) reference, we systematically benchmarked six integration methods *STAligner, SPIRAL, DeepST, GraphST, spatiAlign*, and *PRECAST* across coronal mouse brain dataset (Sample 7) sequenced using “10X_FFPE,” “10X_Normal,” and “10X_DAPI.” Fig. 8 presents three coronal mouse brain slices obtained using distinct experimental protocols. The spatial transcriptomic spots were visualized and colored according to slice indexes, anatomical annotations, clustering results, and spatial domain identification.

**Fig. 8:**
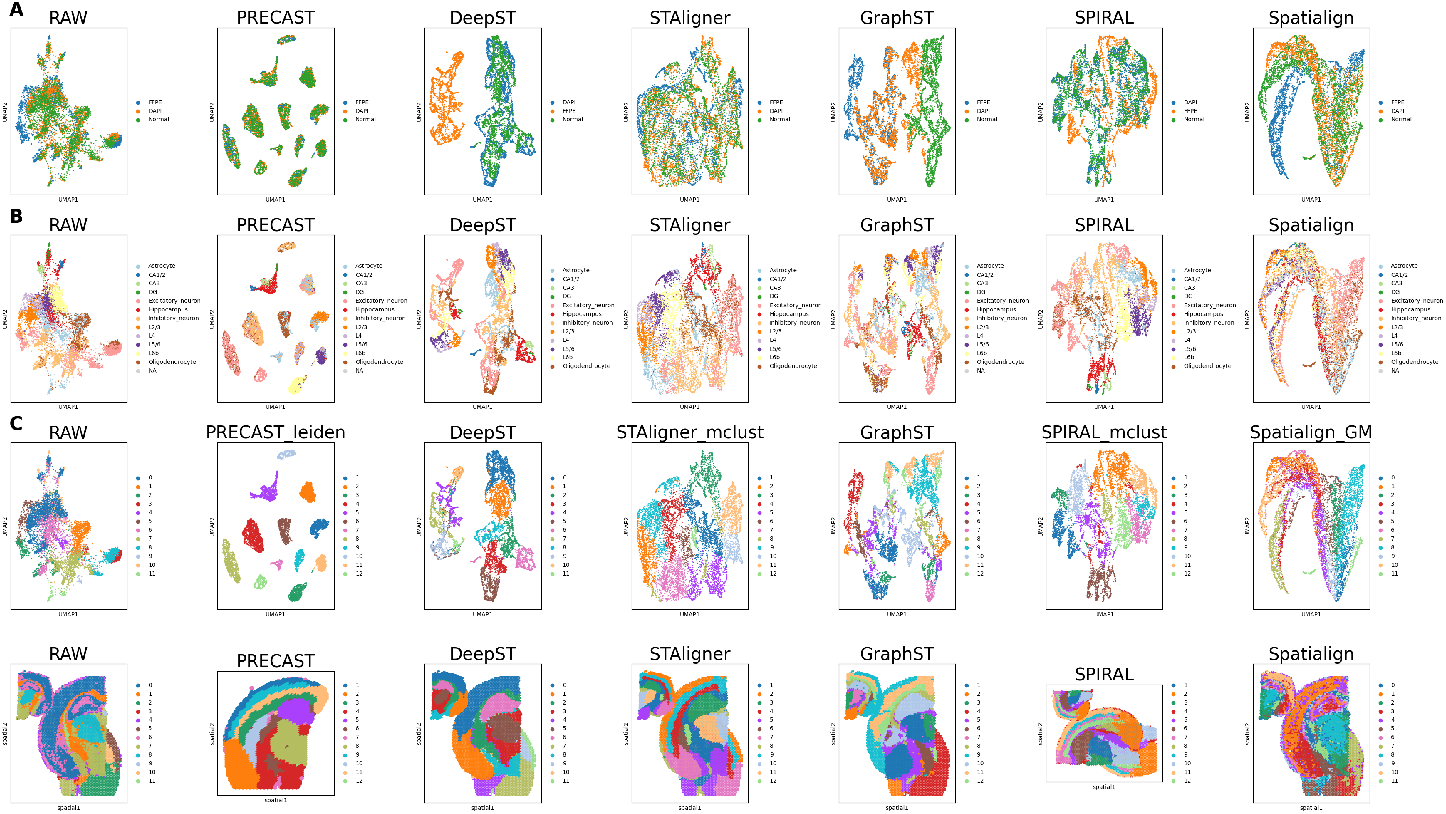
Visualization of the batch effect correction results from six ST methods using the Coronal mouse brain dataset (Sample 7). The dataset includes tissue slices of three protocols: “10X_FFPE,” “10X_Normal,” and “10X_DAPI”. Panels A-C show UMAP plots of uncorrected (RAW) data and six ST methods (*PRECAST, DeepST, STAligner, GraphST, SPIRAL*, and *spatiAlign*). Spots are colored by A: tissue slice indexes, B: manual annotations, and C: clustering outcomes. Panel D comparatively visualizes spatial domain identification performance during cross-protocol integration.

Methods such as *SPIRAL* and *STAligner* produced clustering results that closely resembled the manually annotated individual layers, showing strong performance in both batch effect correction and spatial domain identification. These methods facilitated a clearer visualization of spatial layer distributions, effectively preserving their distinct boundaries. In contrast, *DeepST* was only able to mix slices from fresh frozen tissue, consistent with previous results, suggesting limited success in integrating data across protocols. Similarly, *spatiAlign* performed poorly in batch effect correction, with the spatial domains poorly resolved.

Quantitative metrics (Fig. 9) further supported these observations. *SPIRAL* achieved the highest graph connectivity score (0.99) and a strong ASW score (0.87), demonstrating excellent clustering quality. *PRECAST* also performed well, showing good iLISI scores and clustering quality. While *STAligner* effectively removed batch effects, it exhibited slightly lower clustering quality and local similarity compared to *SPIRAL* and *PRECAST*. Meanwhile, *DeepST, GraphST, and spatiAlign* showed bad batch effect correction, with low kBET scores indicating strong residual batch effects. This was visually confirmed in Fig. 8A,C, where the slices remained poorly mixed and the clustering was more chaotic. with low kBET scores indicating strong residual batch effects. *DeepST* also scored poorly on iLISI (0.07) and showed only moderate ASW.

**Fig. 9:**
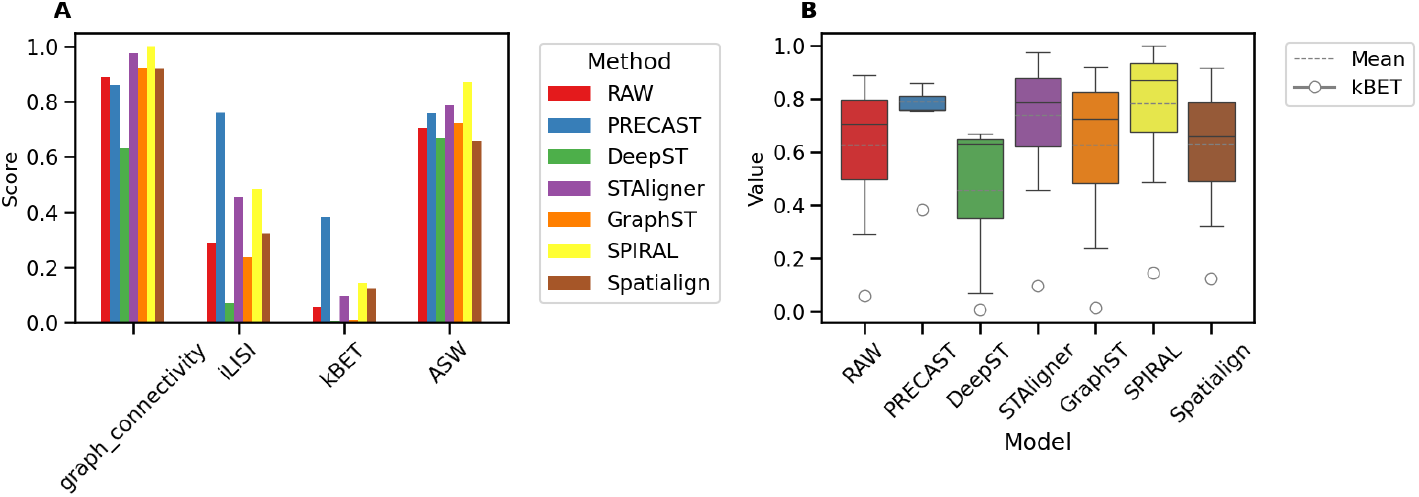
The figure compares six methods using Coronal mouse brain (Sample 7) based on metrics related to batch effect correction in spatial transcriptomics. A: presents the scores of each method on four metrics (GC, iLISI, kBET, ASW); B shows the mean values and quartile distributions of three metrics, (GC, iLISI, and ASW), and kBET in circle.

##### Cross-platform Integration

Due to the lack of available layer information, we could only visualize batch mixing, clustering results, and spatial domains in Fig. 10. The spots were colored based on slice indexes (Fig. 10A) and clustering results (Fig. 10B). Fig. 10C illustrated spatially resolved tissue domains through projection mapping. Among the tested methods, *STitch3D* failed to integrate data across the three platforms due to its reliance on *PASTE* which is incompatible with Stereo-seq and lacks batch effect correction. Similarly, *PRECAST*, designed for single-platform datasets, struggled with crossplatform integration. Based on previous studies, we found that *DeepST* can integrate data from 10X and Stereo-seq, however, due to memory constraints, *DeepST* was excluded from our study. Consequently, we focused on evaluating the remaining methods—*SPIRAL, STAligner, GraphST*, and *spatiAlign*—using qualitative UMAP visualizations and quantitative metrics.

**Fig. 10:**
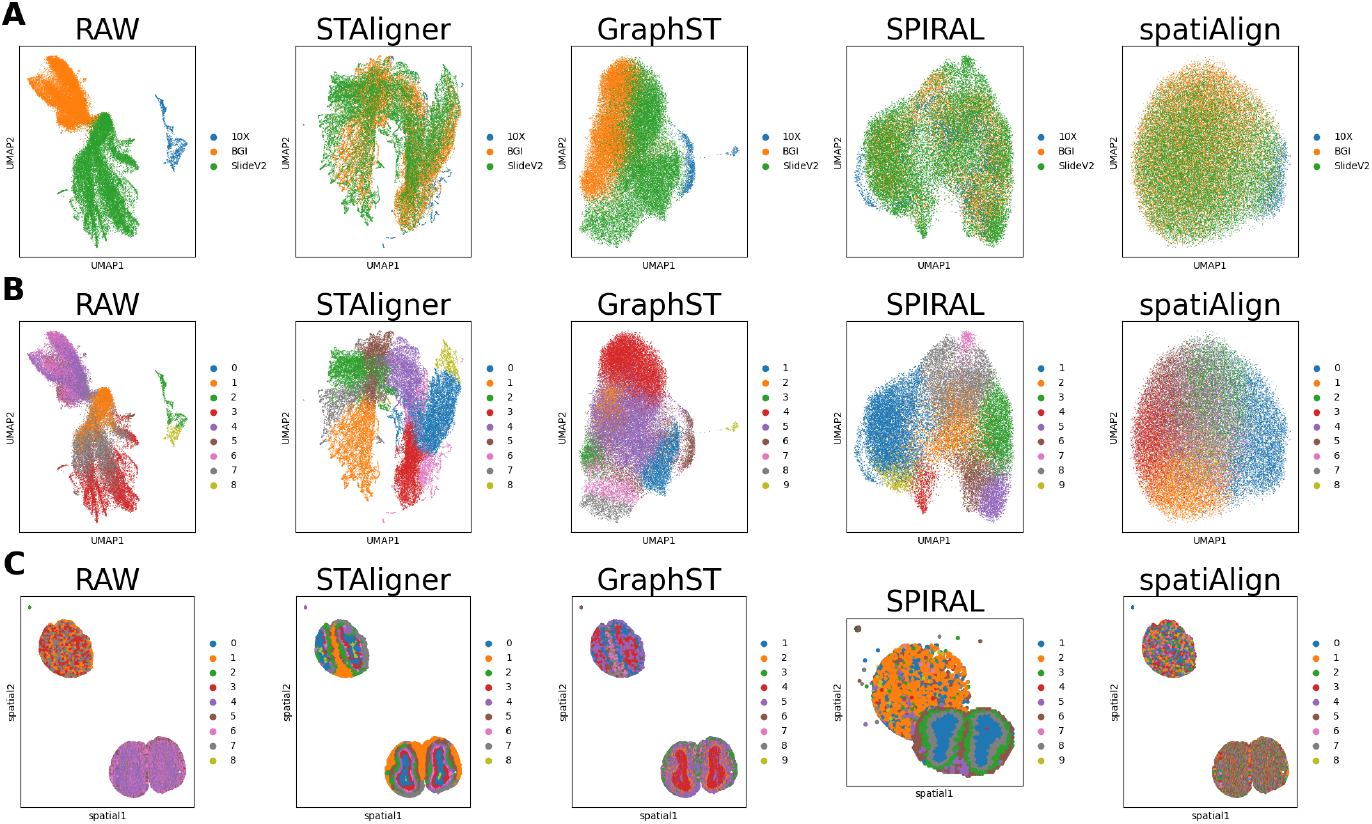
Visualization of the batch effect correction results from four ST methods using the Mouse olfactory bulbs (OB) dataset (Sample 8). The dataset contains tissue slices from three platforms: “10x Visium”, “Stereo-seq”, and “Slide-seq V2”. Panels A-B display UMAP plots of uncorrected data (RAW) and four ST methods (*STAligner, GraphST, SPIRAL*, and *spatiAlign*). Spots are colored by tissue slice indexes and clustering results, respectively. Panel C comparatively visualizes spatial domain identification performance during cross-platform integration.

The UMAP visualizations (Fig. 10) revealed that *SPIRAL* and *STAligner* effectively mitigated batch effects, as evidenced by the well-mixed clusters across datasets from the three platforms. Among these methods, *STAligner* exhibited the clearest spatial distribution, with well-preserved organizational structures, exhibiting the most effective batch effect correction. In contrast, *SPIRAL* showed partial confusion in spatial structure, suggesting that while it was effective in reducing batch effects, it did not fully preserve spatial integrity. *spatiAlign*, however, appeared to suffer from over correction, leading to a loss of biological heterogeneity. This was evident in the complete disappearance of slice boundaries and a disordered spatial structure. *GraphST* performed the worst, as the batches remained clustered rather than being evenly mixed, indicating its inability to effectively correct batch effects. From Fig. 10C, it was clear that the slices were not well-aligned. A key reason for this misalignment may be the differences in resolution between platforms, such as the spatial point density of 10X, BGI, and SlideV2.

For quantitative evaluation (Fig. 11), *SPIRAL* achieved the highest iLISI score, indicating effective batch integration, which was also visible in the UMAP plot (Fig. 10C) with well-mixed clusters. In contrast, *GraphST* had the lowest (0.05), reflecting poor batch effects mitigation. *STAligner* achieved a perfect score for graph connectivity (1.00), effectively preserving biological relationships and eliminating batch effects. For kBET, *STAligner* once again outperformed all other methods, showing optimal dataset integration, while the raw data exhibited the poorest performance in both the metrics and UMAP visualization. These results demonstrated that methods like *SPIRAL* and *STAligner* effectively reduce batch effects and preserve clustering, while *GraphST* struggles with batch integration.

**Fig. 11:**
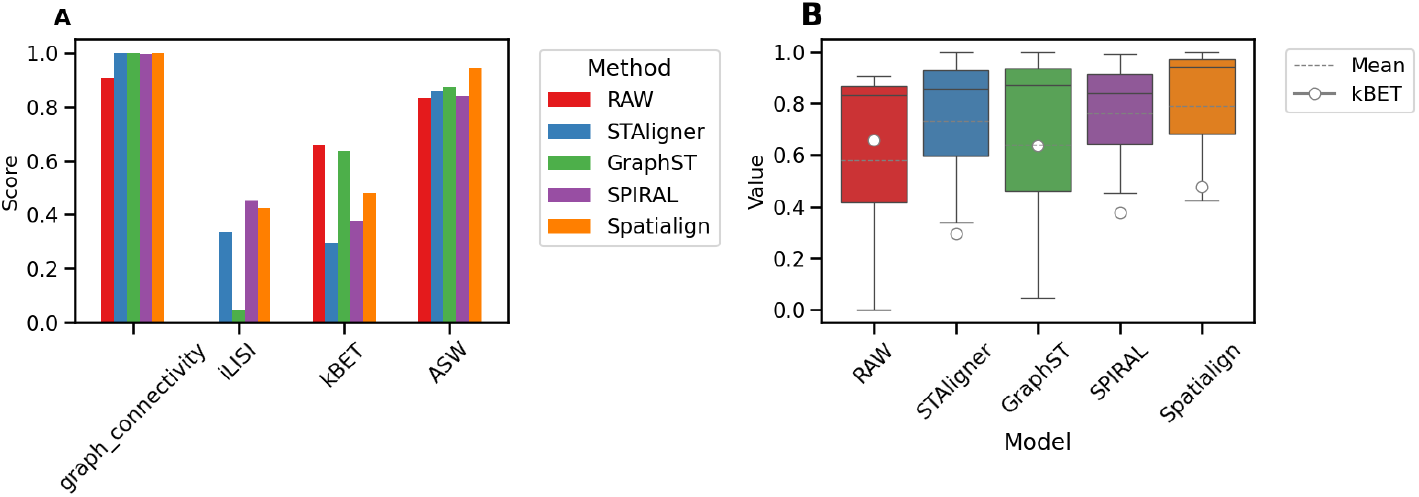
The figure compares four methods using Mouse OB (Sample 8) based on metrics related to batch effect correction in spatial transcriptomics. A presents the scores of each method on four metrics (GC, iLISI, kBET, ASW); B shows the mean values and quartile distributions of three metrics (GC, iLISI, and ASW), and kBET in circle.

In summary, *SPIRAL* demonstrated the highest effectiveness in batch effect correction across various experimental protocols and technological platforms, excelling in both data integration and spatial domain recognition. *STAligner* also performed well, although its clustering quality was slightly lower than that of *SPIRAL. spatiAlign* was designed to effectively integrate data from multiple platforms, leveraging shared information to align batches. *GraphST* struggled significantly with both integration and batch effect correction, making it unsuitable for cross-platform analyses. In addition, *DeepST* was effective only for integrating slices from fresh frozen tissues but failed with datasets involving different protocols. These findings highlighted the importance of carefully selecting methods for spatial transcriptomics analysis based on platforms, experimental protocols, and specific analysis objectives.

### 2.3 Comprehensive comparison

We further evaluated the performance of batch effect correction by calculating the average scores of four metrics—graph connectivity, iLISI, kBET, and ASW—for each of the seven methods across all definitions. The average scores of these four metrics for the methods are summarized in Supplementary file (Table S1). As shown in Fig. 12, in terms of graph connectivity (*p <* 0.01), *STAligner* performs the best, with a mean value approaching 1.00, indicating more consistent layers between batches and more stable clustering. iLISI (*p <* 0.01) shows the highest score for *PRECAST*, with most of its layers coming from the same batch, suggesting smaller batch effects and better separation between batches. For kBET (*p <* 0.05), *PRECAST* achieves the highest scores, as most of the layers within the same batch are well-mixed, indicating minimal batch effects. The ASW (*p <* 0.01) scores of *STitch3D, SPIRAL*, and *GraphST* are all above 0.90, highlighting higher tightness and separation between layers with smaller batch effects. Overall, *spatiAlign* has low scores across each metric, reflecting a bad ability to remove batch effects.

**Fig. 12:**
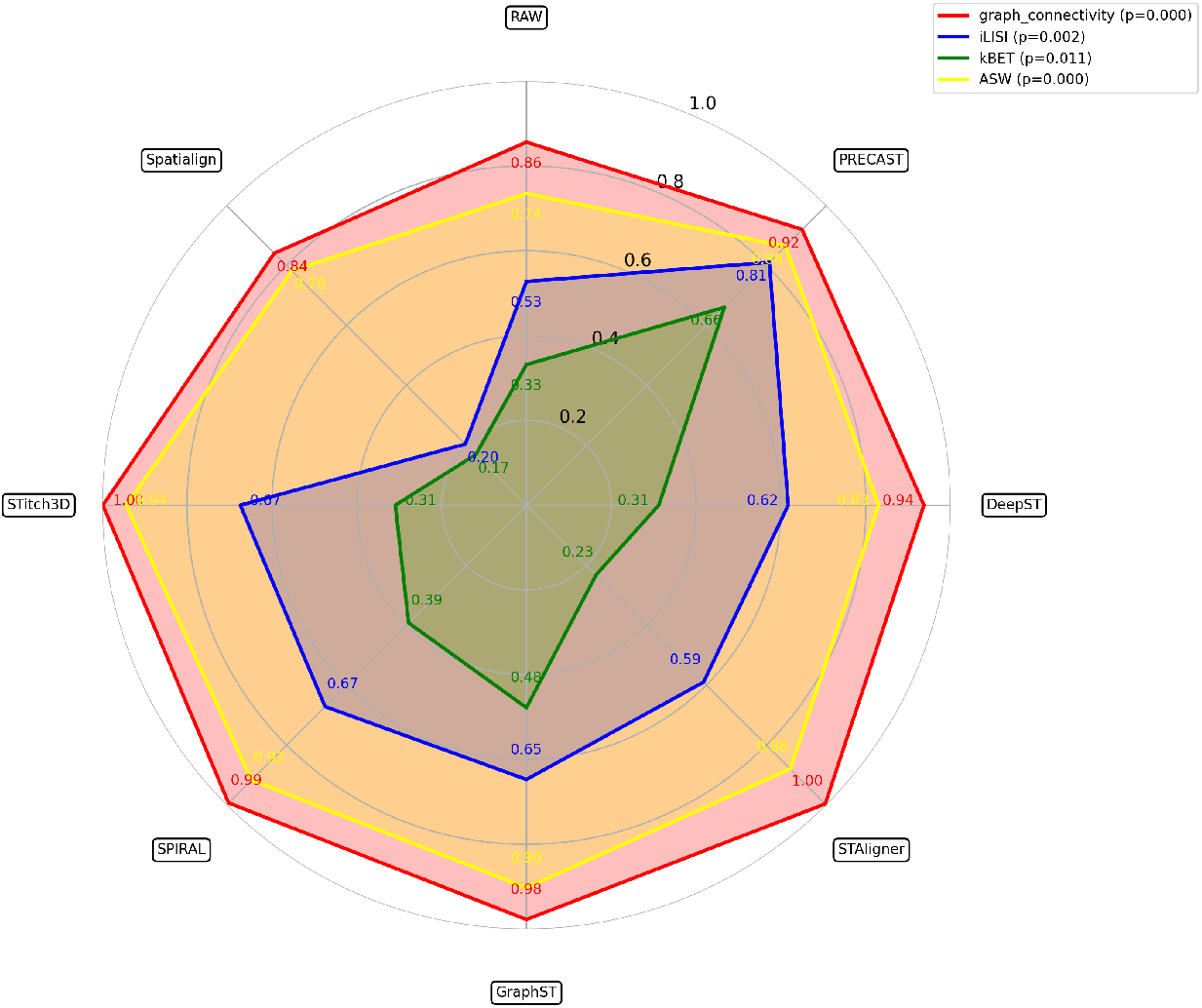
Comparison of the performance evaluation of spatial transcriptomics batch correction algorithms (*STitch3D, Spiral, GraphST, Precast, DeepST, STAligner*) across four key metrics: graph connectivity, iLISI, kBET, and ASW. Radar axes are scaled from 0 (the worst) to 1 (the best), with filled areas indicating algorithm performance profiles.

### 2.4 Comparison of spatial transcriptomic analysis pre- and post-batch effect correction

In this section, we selected *STAligner* for downstream analysis due to its well applicability across all definitions in our study, its ability to preserve spatial geometry information, and its lower system memory requirements. Based on these advantages, we applied *STAligner* to the 12 slices of the DLPFC dataset and evaluated its performance against the uncorrected data in various aspects, including slice integration, batch effect correction, clustering, spatial domain identification, trajectory inference, and differentially expressed gene (DEG) analysis. Additionally, we evaluated batch effect metrics to assess performance, as shown in the Fig. 13.

**Fig. 13:**
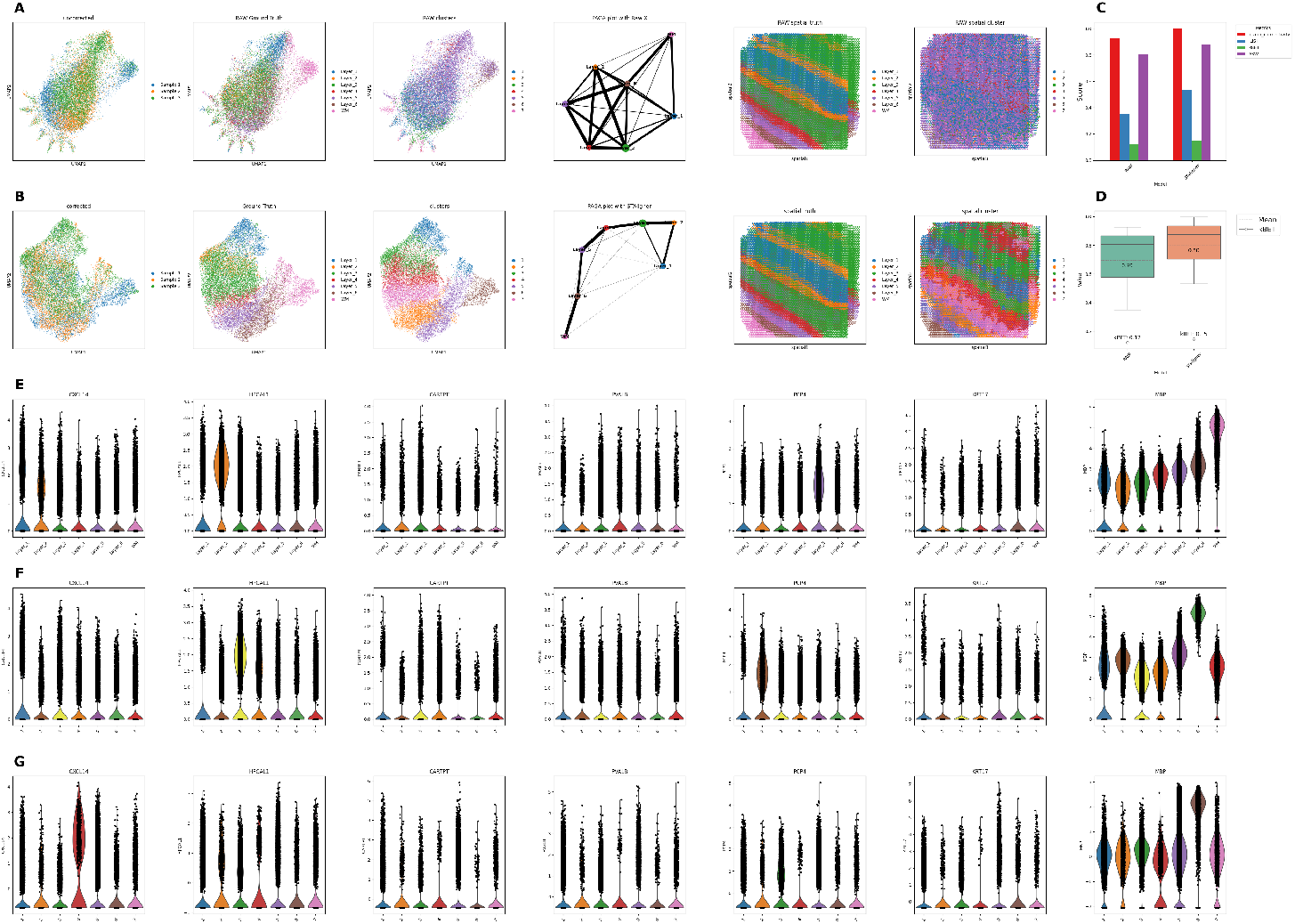
Panels A-G comprehensively evaluate the performance of *STAligner* against uncorrected (RAW) data. A-B: Integration of 12 slices visualized by UMAP embeddings (left: colored by sample index, manual annotation, and clustering) and spatial analyses (right: trajectory inference and domain identification), where A represents RAW data (without batch correction) and B represents *STAligner*-corrected data. C: Summary of four evaluation metrics (GC, iLISI, kBET, ASW). D: Distribution of metrics: boxplots for GC/iLISI/ASW (median and quartiles), circle plot for kBET. E-G: Violin plots show layer-specific marker gene distributions, where dots represent individual spots and black bars indicate median expression. Panels correspond to (E) ground-truth tissue layers in *STAligner*-corrected data, (F) spatial domains identified via *STAligner* clustering, and (G) clusters from RAW data.

*STAligner* significantly improved slice integration, as demonstrated by more cohesive clusters and clearer separation of layers. The UMAP before (Fig. 13A) and after (Fig. 13B) correction show a substantial reduction in batch effects, while preserving the biological structure. In particular, the PAGA plots reveal that *STAligner* captures a more coherent trajectory structure after correction, providing a robust representation of underlying biological processes. The spatial domain of the corrected data align more closely with the ground truth layers. While both the raw and corrected data show some alignment, *STAligner* achieves more distinct and homogeneous spatial domains. This was further supported by higher graph connectivity, iLISI, and ASW scores (Fig. 13C-D), indicating that *STAligner* performs more consistently across different slices. By preserving the biological signals while eliminating the batch effects, *STAligner* establishes a solid foundation for downstream spatial transcriptomics analyses.

To evaluate the impact of batch effect correction on gene expression analysis, we conducted three types of differentially expressed gene (DEG) analyses: (i) reference markers based on known biological expectations, (ii) markers identified from *STAligner*-processed data using its clustering results, and (iii) markers identified from the uncorrected integrated raw data, which served as the control.

In terms of spatial domain delineation and gene expression visualization (e.g., PCP4, HPCAL1, MBP), the corrected data revealed clearer layer-specific expression patterns, better aligning with biological expectations (Fig. 13E). In contrast, the uncorrected data exhibited minimal variability between layers, likely due to residual batch effects masking true biological differences (Fig. 13F-G). For instance, UMAP visualization (Fig. 13B) initially suggested that cluster 3 corresponded to `layer2` and cluster 2 to `layer5`, which was further supported by differential gene expression analysis. Specifically, the known `layer2` marker gene HPCAL1 was predominantly expressed in cluster 3 (*p <* 0.001), while the `layer5` marker gene PCP4 was specifically enriched in cluster 2 (*p <* 0.001; Fig. 13F). Without batch correction, these spatial expression patterns were obscured (Fig. 13G), highlighting the necessity of batch effect correction in reducing confounding variability and enabling the identification of biologically meaningful marker genes in spatial transcriptomics.

## 3. Discussion

### 3.1 Batch effect correction

Batch effects refer to systematic differences caused by non-biological factors such as experimental conditions, operators, reagent batches, equipment status, and environmental conditions. These differences can significantly impact experimental outcomes, even in the absence of true biological variation. In spatial transcriptomics, understanding the characteristics of the dataset—such as spatial resolution, tissue complexity, and platform variability—is crucial, as these factors influence the manifestation of batch effects. Optimizing analytical parameters such as spatial smoothing and distance metrics is essential for minimizing batch effects and improving data integration.

Addressing batch effects is essential for ensuring data quality, accurate biological interpretation, and reliable downstream analyses. With recent advancements in spatial transcriptomics, batch correction has become increasingly important. For single-sample analysis, batch correction improves 3D spatial reconstruction and gene expression accuracy[13][32]. In multiple-sample studies, it facilitates data integration, enabling meaningful inter-sample comparisons, which are particularly important in clinical research[45]. Moreover, batch correction has been shown to enhance machine learning model performance, leading to more precise predictions of biological outcomes[24]. When integrating datasets derived from different technology platforms, effective batch correction mitigates platform-specific biases, enhancing comparability and enabling deeper biological insights.

Traditional workflows involve several steps, including initial assessment, normalization, batch effect correction, and quality control[46]. Common normalization techniques, such as Quantile Normalization and Z-score normalization, are often applied to standardize gene expression data and reduce technical variation. Widely used methods such as ComBat[47], Mutual Nearest Neighbors (MNN)[48], Harmony[49], and *Seurat*[50] have been effective in single-cell transcriptomics[51]. Some of these methods have also been applied to spatial transcriptomics workflows. However, these methods often fail to maintain spatial integrity. While these methods correct expression-level batch effects, they may disrupt the spatial relationships between tissue structures and cellular microenvironments, which are crucial in spatial transcriptomics.

#### 3.1.1 Method comparison and suggestion

In this study, we systematically evaluated seven representative ST methods using four batch correction metrics: kBET, ASW, iLISI, and GC. These metrics, initially designed for single-cell transcriptomics, have been adapted for spatial transcriptomics due to inherent similarities in data structure and batch effect characteristics. Beyond their applicability, they play a crucial role in assessing batch mixing in lower-dimensional embedded space, which is essential for evaluating the effectiveness of batch correction in spatial transcriptomics.

Among the four metrics, kBET consistently yields lower scores compared to ASW, iLISI, and GC. This difference arises from kBET’s stringent chi-square-based testing of local batch proportion equilibrium. In contrast, other metrics tolerate batch proportion deviations through distinct mechanisms: iLISI quantifies batch mixing diversity via inverse Simpson index within local neighborhoods, ASW leverages silhouette widths to assess global batch mixing while preserving biological alignment, and GC assesses preservation of biologically coherent layer-layer neighborhood graphs.

To systematically evaluate batch correction performance, we integrated multiple perspectives by combining UMAP visualizations, spatial projections, and four quantitative metrics. Through this holistic analysis, we generated a suggestion table (Table 1) to guide method selection for spatial trascriptomics studies. This table consolidates both quantitative metric scores and qualitative insights from visual inspections. Our findings emphasize that alignment and integration methods play a crucial role in mitigating batch effects, particularly in multi-slice spatial transcriptomics datasets.

Among the methods assessed, *STAligner* demonstrates superior integration, with clearer separation between spatial domains and stronger connectivity. This can be attributed to its use of triplet learning and spatially aware autoencoders, which likely help reduce batch effects. *SPIRAL* also performs well, particularly in protocol/platform integration, showing good separability across batches both in space and gene expression. It excels in data integration and spatial domain recognition, potentially due to its combination of GraphSAGE with domain adaptation and the use of Gromov-Wasserstein distance for spatial alignment, which may improve data consistency. *GraphST* performs well in batch effect correction, particularly for inter-slice and inter-sample analysis, which may be attributed to its unique approach. Instead of explicitly removing batch factors, it smooths feature distributions across batches through iterative aggregation of neighboring spot representations and graph self-supervised contrastive learning. This dual mechanism likely ensures spatial preservation while mitigating batch effects, which could explain its outperformance compared to simpler correction strategies. However, *GraphST* is limited by its reliance on accurate coordinate alignment, restricting its use for diverse datasets. It also struggles with batch effects when integrating data from multiple platforms. *PRE-CAST* performs well in certain datasets but struggles with preserving spatial geometry information, potentially leading to more noticeable segregation in latent space. It employs a projection strategy that accounts for shifts in cluster centroids during dimension reduction, which might implicitly correct batch effects. By utilizing the intrinsic Component Adjusted Regression (CAR) method, *PRECAST* ensures smoothness across neighboring embeddings while adjusting for shifts between batches, helping to maintain spatial integrity and improving data integration. *DeepST* also has limitations. As a deep learning-based approach, it can be computationally expensive for large datasets, particularly when handling varying dataset resolutions and mitigating batch effects across slices.

### 3.2 Key Challenges in batch effect correction of spatial transcriptomics

Correcting batch effects in spatial transcriptomics presents several challenges arising from both technical and biological factors across four key categories. For tissue slices derived from the same sample, variations in spatial resolution and imaging quality across slices often introduce technical biases, making it difficult to maintain 3D spatial continuity after correction. When analyzing multiple samples, biological differences are confounded with technical biases, while variations in sample preparation and experimental protocols further complicate normalization. Normalizing data across samples while preserving sample-specific biological signals remains a key challenge. Additionally, integrating sequencing-based and imaging-based approaches, which rely on fundamentally different measurement principles, poses significant difficulties. For tissue slices generated using different technological platforms, differences in resolution, data sparsity, and platform-specific biases create additional barriers. Within the same tissue slice, intrinsic tissue heterogeneity further complicates the distinction between biological variation and technical artifacts. Moreover, issues such as uneven staining, spatial artifacts, and edge effects can distort spatial gene expression patterns. These factors highlight the complexity of batch effect correction in spatial transcriptomics and the need for robust methods that preserve both spatial and biological integrity.

Another critical challenge in this context is the risk of over-correction of batch effects. Excessive correction can lead to the loss of important biological and spatial information. Over-correction may blur tissue boundaries and disrupt spatial gradients, making it difficult to distinguish regions and misalign biological clusters. It is essential that batch correction methods preserve both spatial continuity and biological clustering. Over-correction can lead to inaccurate or less informative results by merging distinct clusters and disrupting manual annotations. These concerns underscore the need for methods that not only address batch effects but also ensure that biological insights are not compromised in the process.

Additionally, computational cost remains a major challenge, particularly for large, high-resolution spatial transcriptomics datasets. Many methods scale poorly with increasing data size, limiting their applicability in real-world scenarios. While the integration of spatial transcriptomics data across platforms holds immense potential for advancing tissue biology and disease research, challenges such as persistent batch effects and data heterogeneity remain unresolved. Future efforts should focus on developing robust, scalable methods, promoting data sharing and standardization protocols. By addressing these challenges, the field of spatial transcriptomics can continue to grow as a transformative tool for understanding complex biological systems and diseases.

### 3.3. Limitations of our study

While numerous spatial transcriptomics approaches incorporate batch effect correction, our selection of seven widely utilized methods was guided by specific considerations. Initially, thirteen methods were identified, but many lacked clear batch effect correction processes or sufficient detail for effective evaluation. The seven selected methods are widely adopted, well-documented, and supported by available tools, making them practical and reliable choices for comparison. This selection ensures meaningful results, providing researchers with accessible and well-validated correction strategies. Moreover, biological validation of corrected data is difficult, especially in cases where functional regions are poorly annotated or lack well-defined biological ground truth.

While our work defines batch effects in spatial transcriptomics and evaluates existing methods, further methodological innovation is needed to improve scalability, develop biologically informed validation strategies, and establish robust evaluation criteria suited to the complexity of spatial transcriptomic datasets.

## 4 Conclusion

In this study, we defined batch effects in spatial transcriptomics, reviewed methods for their correction, and evaluated seven widely-used approaches across datasets from the DLPFC, human breast cancer, mouse olfactory bulb, and coronal mouse brain. Our analysis revealed that no single method outperformed all others, but specific methods showed their strengths under certain conditions. Among the methods, *STAligner, SPIRAL, GraphST*, and *PRECAST* performed well with datasets from the same platform. *STAligner* demonstrated stronger connectivity between slices, SPIRAL effectively aligned slices and controlled batch effects removal, *PRECAST* performed well in integrating slices from different samples and protocols. Both *SPIRAL* and *STAligner* were effective in recognizing spatial domains across different protocols and platforms, showing robust batch mixing. Overall, these findings highlight the importance of selecting appropriate methods based on specific dataset characteristics and study objectives to achieve optimal batch effect correction in spatial transcriptomics.

## 5 Methods

Table 2 summarizes key methods with inherent batch effect correction in spatial transcriptomics, categorizing them based on their approach and highlighting their key techniques and outputs.

**Table 2:**
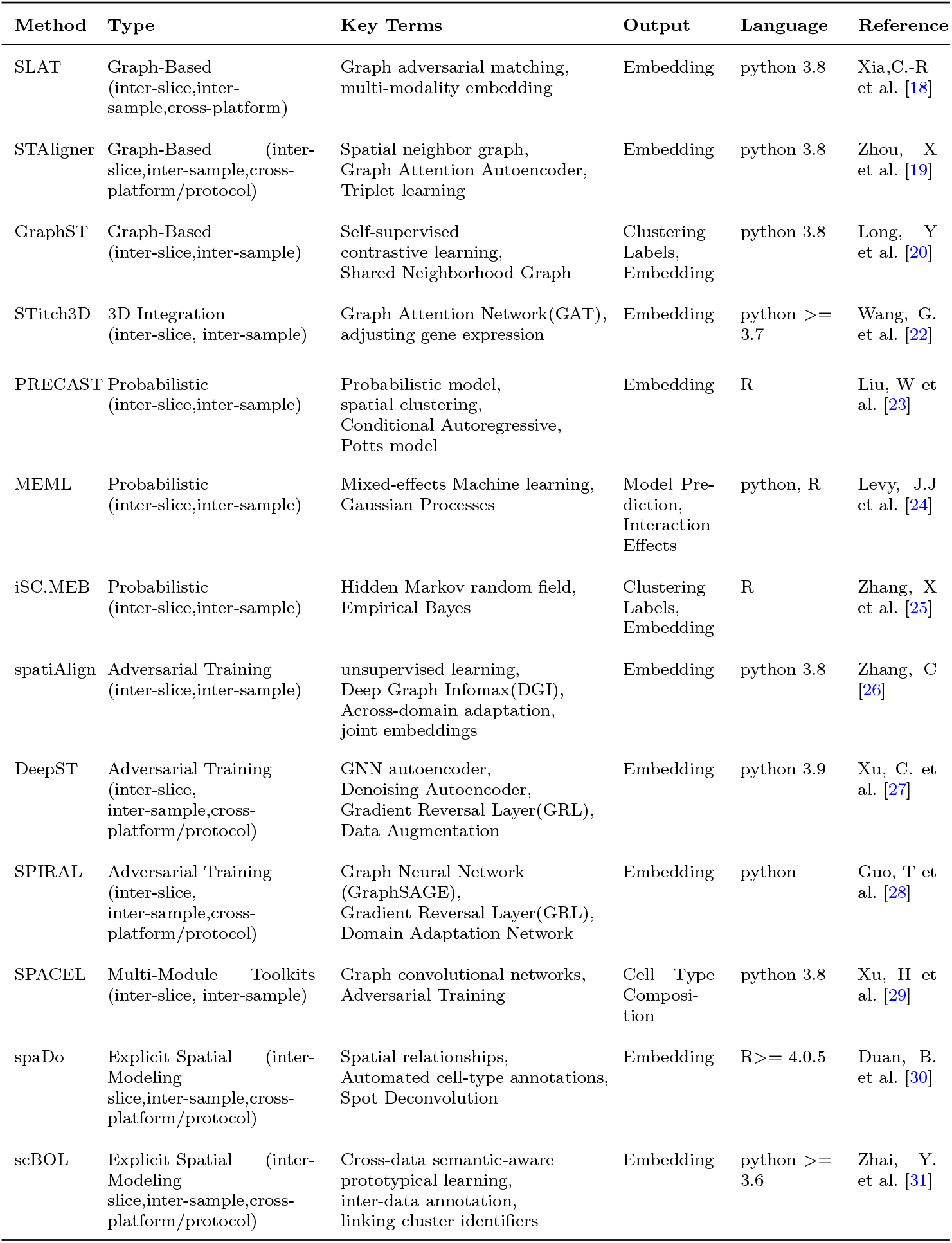
Comparison of methods with inherent batch effect correction.

Graph-based methods like *SLAT, STAligner*, and *GraphST* utilize concepts such as graph adversarial matching and spatial neighbor graphs for embedding and clustering. *STitch3D* removes batch effects by adjusting gene expression and accounting for slice- and gene-specific effects while integrating multiple spatial transcriptomics slices into a unified 3D representation. Probabilistic methods like *PRECAST, MEML*, and *ISC*.*MEB* focus on spatial clustering and across-domain adaptation for generating embeddings and predictions. *DeepST, spatiAlign*, and *SPIRAL* apply adversarial training and domain adaptation, incorporating graph neural networks for batch effect correction. *SPACEL* uses graph convolutional networks and adversarial training for cell type composition estimation, while spatial modeling methods like *spaDo* and *scBOL* focus on spatial relationships and automated annotations. This overview highlights the methods’ diverse approaches to batch effect correction, spatial domain recognition, and data integration.

### 5.1 Methods with inherent batch effect correction

Thirteen methods have been developed to address the challenges of batch effects and integrate spatial transcriptomics data from different slices, samples, and technology platforms. Below, we present an overview of key methods used in spatial transcriptomics integration:

#### 5.1.1 SLAT (Spatially-Linked Alignment Tool)

*SLAT*[18] is a tool designed to align slices from the same sample, across different samples, and between different technological platforms/protocols. The method employs a graph adversarial matching strategy to align data in a shared low-dimensional space using Singular Value Decomposition (SVD) during homogeneous alignment, effectively mitigating batch effects between datasets. For cross-modality spatial alignment, such as integrating Spatial-ATAC-seq and Stereo-seq data, *SLAT* uses a graph-linked multi-modality embedding strategy to project data from different modalities into a common embedding space, addressing batch effects and preserving both modality-specific and spatial information.

#### 5.1.2 STAligner (Graph Attention Neural Network for ST Data)

*STAligner*[19], built on the STAGATE model, is an advanced integration method for spatial transcriptomics (ST) data that aligns slices from the same sample, different samples, and different technology platforms/protocols. It constructs a spatial neighbor graph based on spot coordinates and uses a graph attention autoencoder to create spatially-aware embeddings. The method incorporates spot triplet learning based on mutual nearest neighbor (MNN) principles to minimize the distance between anchor-positive pairs while maximizing the distance between anchor-negative pairs during slice alignment. This strategy effectively reduces batch effects in the latent space, enhancing integration quality and spatial coherence.

#### 5.1.3 GraphST (Self-Supervised Contrastive Learning)

*GraphST* [20] introduces the PASTE technique for aligning spatial coordinates to reduce batch effects during horizontal integration. When vertically integrating slices from multiple samples, the method first aligns H&E images and constructs a shared neighborhood graph. Through iterative aggregation of neighbor representations, *GraphST* smooths feature distributions and mitigates batch differences. A self-supervised contrastive learning strategy is then employed to reinforce spot embeddings, ensuring that spatially neighboring spots exhibit similar representations, further reducing batch effects and preserving spatial structures.

#### 5.1.4 STitch3D (3D Cellular Structure Reconstruction)

*STitch3D*[22] is a unified framework designed to integrate multiple ST slices to reconstruct 3D cellular structures. This method focuses on identifying spatial domains and aligns slices through iterative closest point (ICP) or PASTE techniques, converting spatial point coordinates into a common coordinate system. It then constructs a 3D neighbor graph using a Combined Coordinate System (CCS) and generates a shared latent space using a Graph Attention Network (GAT). By accounting for slice- and gene-specific effects, *STitch3D* adjusts gene expression, effectively removing batch effects and integrating spatial information across multiple slices.

#### 5.1.5 PRECAST (Probabilistic embedding, clustering, and alignment for integrating ST)

*PRECAST*[23] is a probabilistic model that integrates slices from multiple samples by directly taking normalized gene expression matrices as input. It aligns and estimates joint embeddings for biological effects across different domains. The method applies a simple projection strategy combined with spatial dimension reduction and spatial clustering. It then utilizes aligned representations, intrinsic CAR components, estimated labels, and Potts models to deal with batch effects in spatial data, ensuring that biological signals are preserved while removing unwanted technical variations.

#### 5.1.6 MEML (Mixed-effects machine learning)

*MEML*[24] addresses batch effect correction by modeling batch effects using Gaussian Processes nested within random intercepts, without including random slopes or considering multiple hierarchical levels.

#### 5.1.7 iSC.MEB (Integrated spatial clustering with hidden Markov random field using empirical Bayes)

*iSC*.*MEB*[25] uses a probabilistic approach with a spatial clustering model HMRF and batch effect correction via Empirical Bayes.

#### 5.1.8 spatiAlign (Unsupervised contrastive learning model)

*spatiAlign*[26] performs self-supervised comparative learning through the Deep Graph Information Maximisation(DGI) technique for integrating slices from multiple samples. This approach uses across-domain adaptation techniques to align joint embeddings, effectively accounting for batch effects across tissue sections. By leveraging the DGI framework, *SpatiAlign* enhances the integration process while preserving the spatial context and minimizing the influence of batch effects on biological interpretations.

#### 5.1.9 DeepST (Deep Learning Framework for ST)

*DeepST*[27] is a customizable deep learning framework that integrates spatial transcriptomics data across multiple batches and technology platforms/protocols. It learns joint embeddings across batches and maps them into a shared latent space for integration. The method combines a graph neural network (GNN) autoencoder with a denoising autoencoder to generate a latent representation of augmented ST data. Additionally, *DeepST* introduces a gradient reversal layer (GRL) and domain discriminator through domain adversarial neural networks (DAN), effectively eliminating batch effects and adapting the model to cross-domain differences in data.

#### 5.1.10 SPIRAL (Integrating and aligning ST data)

*SPIRAL*[28] consists of two consecutive modules: spiral-integration and spiral-alignment. The spiral-integration module integrates slices from different samples and technology platforms/protocols, correcting for batch effects by combining an inductive graph neural network (GraphSAGE) with a domain adaptation network. By using the GraphSAGE encoder and performing gradient inversion in a decoupled network, *SPIRAL* effectively removes batch effects during data integration, ensuring that the integrated spatial transcriptomics data maintains biological coherence.

#### 5.1.11 SPACEL (Comprehensive Toolkit for ST)

*SPACEL*[29] is a comprehensive toolkit for processing spatial transcriptomics data, consisting of three modules: Spoint, Splane, and Scube. The Splane module uses cell-type composition information as input and employs adversarial training within a graph convolutional network (GCN) model to correct batch effects while identifying coherent spatial domains across multiple slices.

#### 5.1.12 spaDo (Multi-slice Spatial Transcriptome Domain Analysis)

*spaDo*[30] reduces batch effects by leveraging spatial relationships between spot, automated cell-type annotation, and spot deconvolution techniques. By focusing on spatial context and ensuring consistent cell-type annotation, spaDo aligns cellular features across batches without the need for explicit batch correction steps, enabling the integration of multiple slices. This approach makes spaDo suitable for analyzing spatial transcriptomics data across different samples, platforms, and protocols.

#### 5.1.13 scBOL (Universal cell type identification framework for single-cell and spatial transcriptomics data)

*scBOL*[31] reduces batch effects by utilizing prototypical learning for cluster alignment and semantic anchors to ensure consistent cell-type annotations across samples, platforms, and protocols. This approach aligns clusters across batches using mutual nearest prototypes and refines cell-type alignment at both the prototype and sample levels, resulting in robust integration and mitigation of batch-related variations.

### 5.2 Methods requiring combination with other algorithms for effective batch correction

#### 5.2.1 STAGATE (Adaptive Graph Attention Auto-Encoder)

*STAGATE[32]*, when integrating slices from the same sample or different platforms/protocols, uses an Adaptive Graph Attention Auto-Encoder (AGAE) to construct a shared latent space. This approach can accurately identify spatial domains but only partially mitigates the differences between slices. Therefore, it needs to be combined with *Harmony* to remove batch effects effectively.

#### 5.2.2 SEDR (Deep Embedded Clustering)

*SEDR*[33] integrates multiple slices from the same sample by using the Deep Embedded Clustering (DEC) method, which combines unsupervised deep autoencoders and graph convolutional autoen-coders to construct a shared latent space. DEC aligns data from different batches in this latent space, which helps reduce batch effects. However, the batch effect correction is not significant on its own, necessitating the use of *Harmony* for more effective batch effect correction.

### 5.3 Methods without batch effect correction capabilities

*BASS*[34], *BayesSpace*[35], and *SpaGCN* [36], which are unable to integrate ST datasets from different batches simultaneously. These methods require the use of additional batch effect correction tools, such as *Harmony*, to effectively remove batch effects during integration.

### 5.4 Evaluation Metrics

#### 5.4.1 kBET (k-Nearest Neighbor batch effects Test)

kBET evaluates local batch mixing by testing whether batch label distributions in kNN neighborhoods align with the global batch distribution. The inputs include corrected embeddings, slice (sample) labels, and clustering labels (or ground truth if manually annotated). A high rejection rate indicates poor batch mixing, while a score of 1 (1 minus rejection rate) represents perfect batch integration. Here, this metric is particularly useful for identifying spatially localized batch effects and ensures that spots from different batches are evenly distributed in local spatial neighborhoods. However, due to the local structure of the embedding space, kBET scores may appear favorable even when batch effects persist. To mitigate this issue, raw data were excluded from comparisons, and kBET was reported separately to avoid biasing the overall performance score in this study.

#### 5.4.2 iLISI (Integration Local Inverse Simpson’s Index)

In spatial transcriptomics, iLISI evaluates batch mixing by assessing the diversity of batch labels in spatial neighborhoods. A kNN graph is constructed using corrected embeddings (which incorporate spatial coordinates and gene expression data after integration), and the diversity of batch labels in each neighborhood is calculated using the Inverse Simpson’s Index. Here, scores range from 0 to 1, where 1 indicates perfect batch mixing (spots from different batches are evenly distributed), and 0 indicates strong batch effects (spots from the same batch cluster together). By focusing on the diversity of batch labels in spatial neighborhoods, iLISI ensures that spatially adjacent spots are well mixed across batches rather than being dominated by a single batch.

#### 5.4.3 ASW (Batch Adjusted Silhouette Width)

Batch Adjusted Silhouette Width (ASW) measures batch mixing within spatially defined layer clusters while preserving cluster integrity. The inputs include corrected embeddings, slice (sample) labels, and clustering labels (or ground truth if manually annotated). For each layer, the silhouette width of batch labels is calculated, which measures the relationship between the distances of spots from the same batch and those from different batches. Here, scores are standardized to range from 0 to 1, where 1 represents ideal batch mixing, and 0 indicates strong batch separation.

#### 5.4.4 Graph Connectivity (GC)

Graph Connectivity metric assesses whether spots of the same layer maintain structural and biological coherence in the k-nearest neighbor (kNN) graph after integration. A kNN graph is constructed using low-dimensional embeddings derived from integrated spatial coordinates and gene expression data, typically with Euclidean or cosine distance metrics. For each layer, the score is calculated as the ratio of the size of the largest connected component (LCC) within the subgraph to the total number of spots belonging to that layer. Here, scores range from 0 to 1, where 1 indicates all layer-specific spots form a fully connected subgraph across batches, reflecting successful batch integration without compromising structural integrity.

## 6 Supplementary information

Supplementary results and Figures S1-S6, Table S1.

## 7 Declarations

### 7.1 Ethics approval and consent to participate

Not applicable

### 7.2 Consent for publication

Not applicable

### 7.3 Availability of data and materials

Our benchmarking codes and details are now publicly available at https://github.com/Yingxinzz/model

### 7.6 Authors’ contributions

Qingzhen Hou conceived and led this work. Yingxin Zhang and Ming Jing drafted the initial manuscript, organizing the content and ensuring logical flow. Ruotong Liu, Na Zhou, and Guoneng Yuan were responsible for data visualization and analysis, processing and analyzing the experimental data to support the study’s results. Fuzhong Xue and Qingzhen Hou supervised the project and provided critical revisions to the manuscript. All authors reviewed and approved the final version of the manuscript.

## 7.7 Acknowledgements

Not applicable

