## Supplementary material for "Towards a Better Understanding of Batch Effects in Spatial Transcriptomics: Definition and Method Evaluation": &#34917;&#20805;&#25991;&#20214;

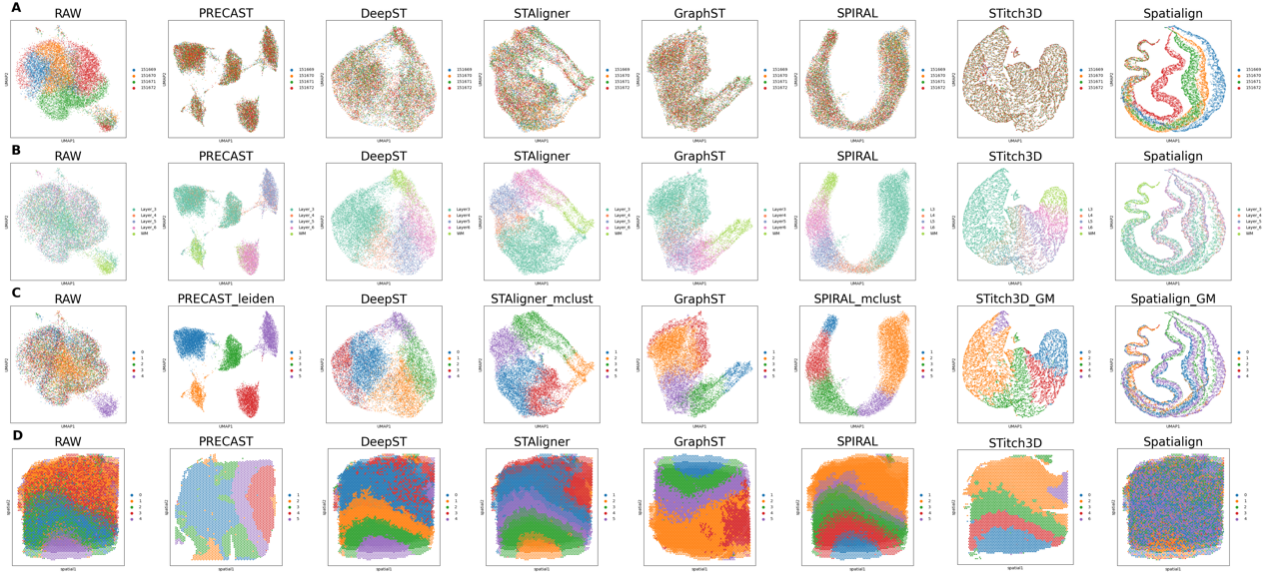

**Figure S1.** Visualization of the batch effects correction results from seven ST methods using DLPFC (Sample 2) dataset. The dataset includes four tissue slices: “sliceA: 1516769”, “sliceB: 151670”, “sliceC: 151671”, and “sliceD: 151672”. Panels A-C show UMAP plots of uncorrected (RAW) data and seven ST methods (PRECAST, DeepST, STAligner, GraphST, SPIRAL, STitch3D, and spatiAlign). Each UMAP plot is colored by three different setups, A: slice indexes, B: manual annotations, and C, clustering results. Panel D uses clustering-label visualizations to compare spatial domain identification performance during non-consecutive slices integration.

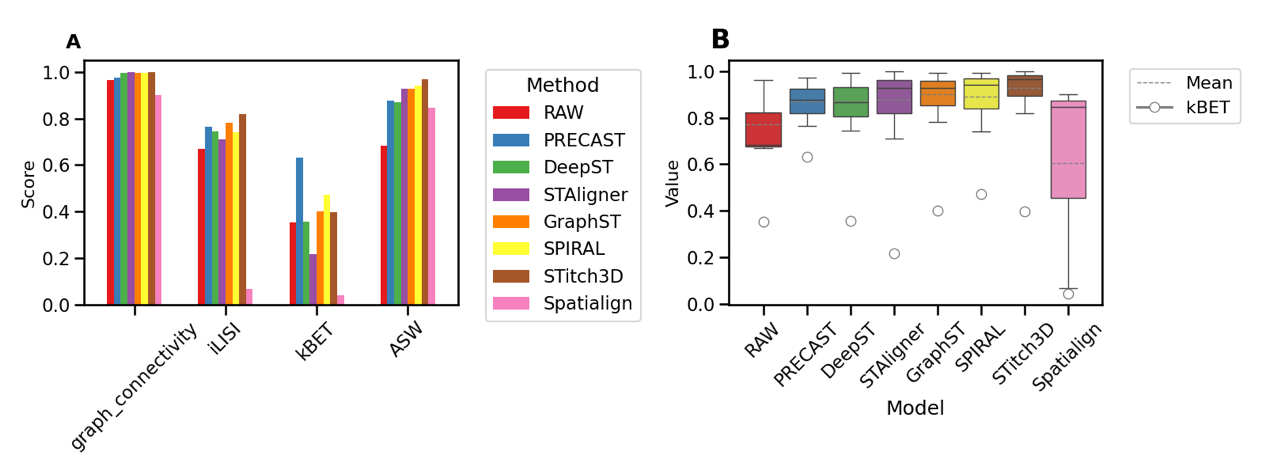

**Figure S2.** The figure compares seven methods using DLPFC (Sample 2) based on metrics related to batch effects correction in spatial transcriptomics. A presents the scores of each method on four metrics (GC, iLISI, kBET, ASW); B shows the mean values and quartile distributions of three metrics, (GC, iLISI, ASW), and kBET in circle.

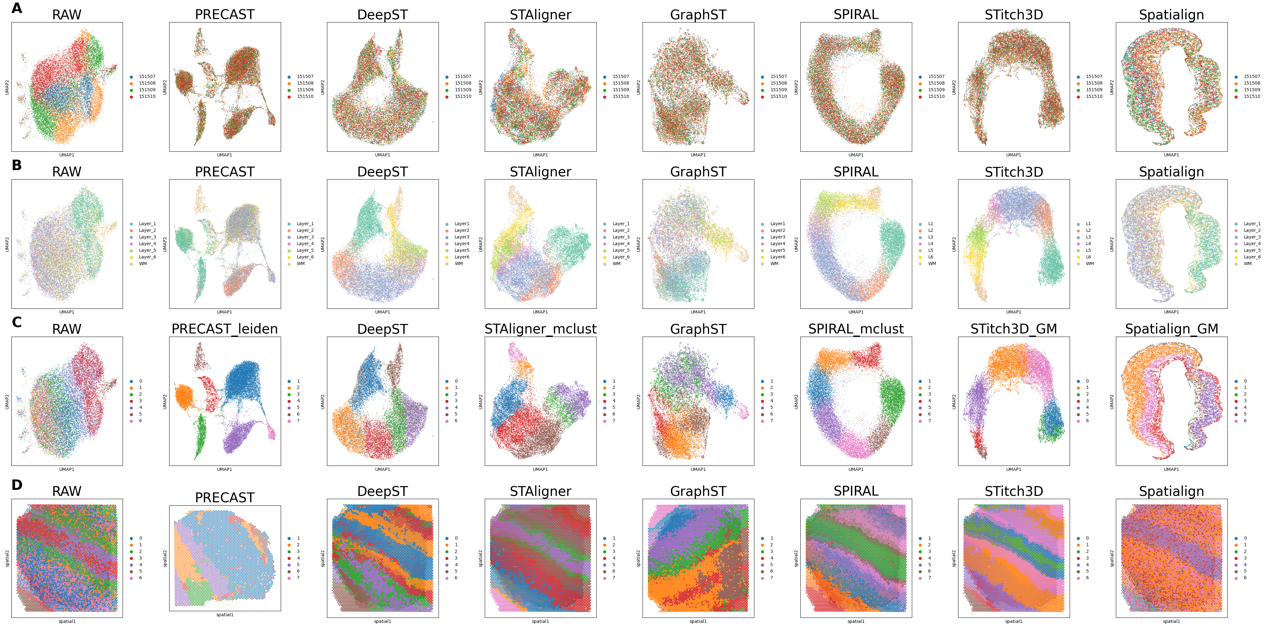

**Figure S3.** Visualization of the batch effects correction results from seven ST methods using DLPFC (Sample 3) dataset. The dataset includes four tissue slices: “sliceA:151507”, “sliceB:151508”, “sliceC:151509”, “sliceD:151510”. Panels A-C show UMAP plots of uncorrected (RAW) data and seven ST methods (PRECAST, DeepST, STAligner, GraphST, SPIRAL, STitch3D, and spatiAlign). Each UMAP plot is colored by three different setups, A: slice indexes, B: manual annotations, and C: clustering results. Panel D uses clustering-label visualizations to compare spatial domain identification performance during non-consecutive slices integration.

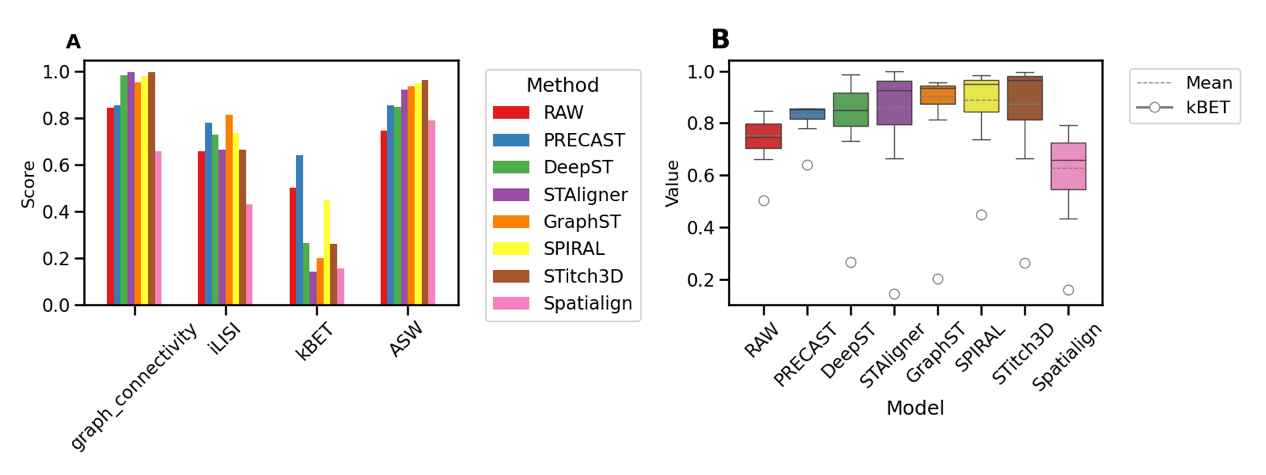

**Figure S4.** The figure compares seven methods using DLPFC (Sample 3) based on metrics related to batch effects correction in spatial transcriptomics. A presents the scores of each method on four metrics (GC, iLISI, kBET, ASW); B shows the mean values and quartile distributions of three metrics, (GC, iLISI, ASW), and kBET in circle.

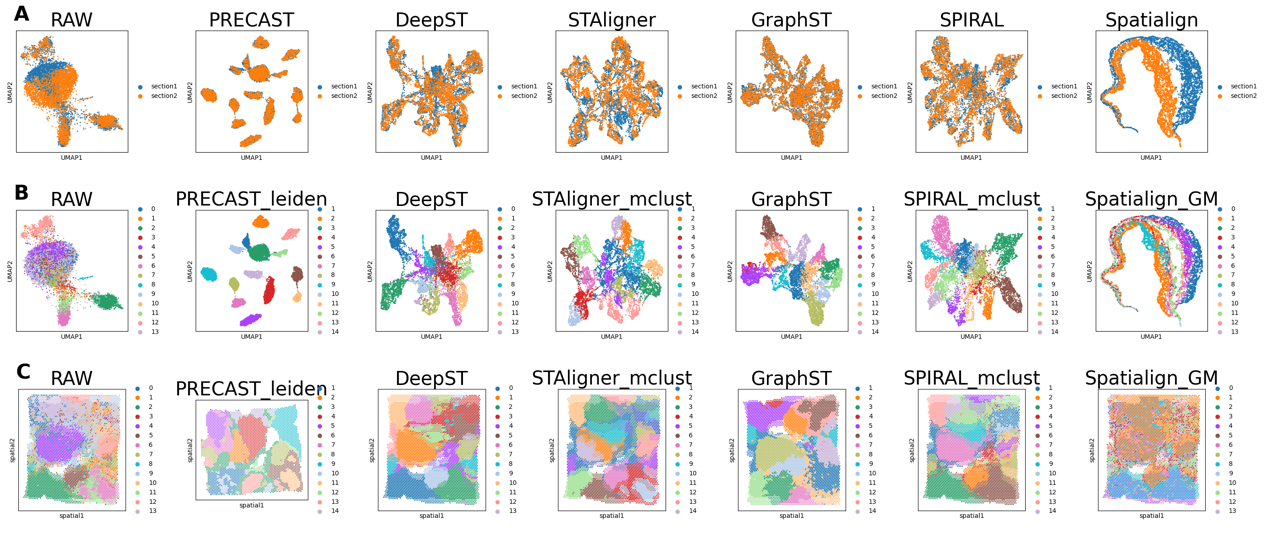

**Figure. S5.** Visualization of the batch effects correction results from six ST methods using Human Breast Cancer (Sample 4) dataset. The dataset includes two tissue slices: “section 1” and “section 2”. Panels A-B show UMAP plots of uncorrected (RAW) data and seven ST methods (PRECAST, DeepST, STAligner, GraphST, SPIRAL, and spatiAlign). Each UMAP plot is colored by two different setups, A: slice indexes, and B: clustering results. Panel C uses clustering-label visualizations to compare spatial domain identification performance during non-consecutive slices integration.

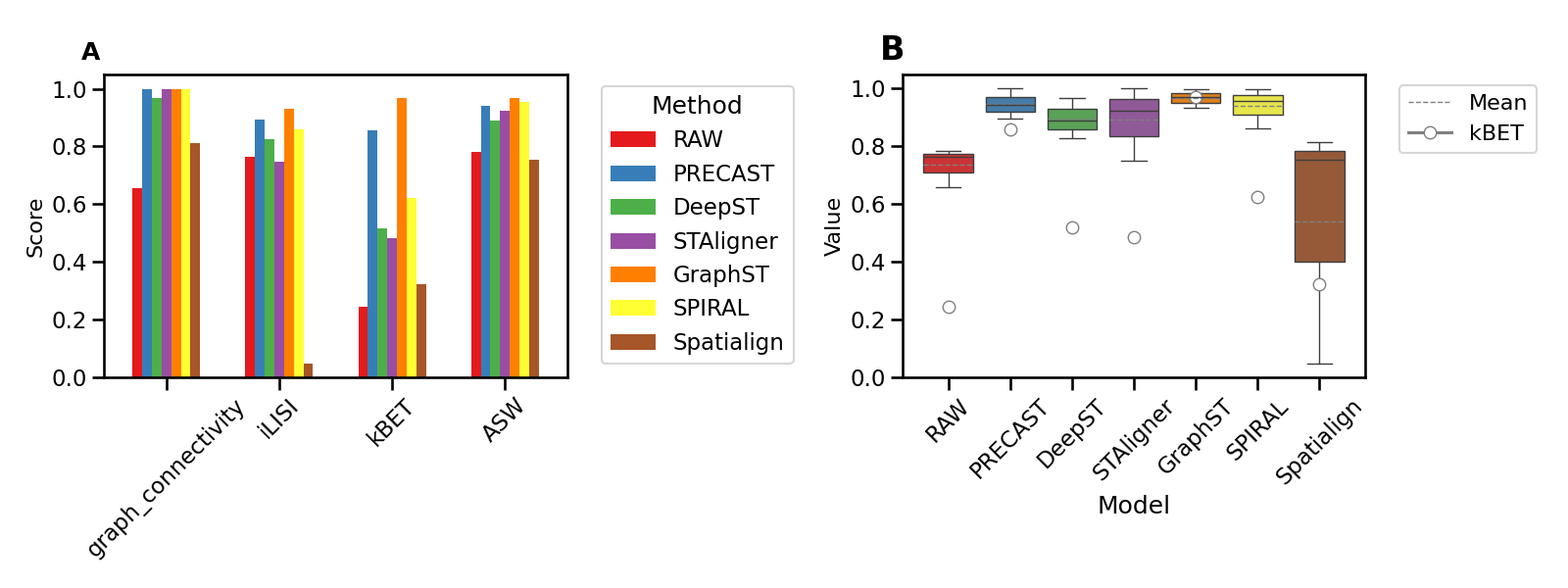

**Figure S6.** The figure compares seven methods using Human Breast Cancer (sample 4) based on metrics related to batch effects correction in spatial transcriptomics. A presents the scores of each method on four metrics (GC, iLISI, kBET, ASW); B shows the mean values and quartile distributions of three metrics, (GC, iLISI, ASW), and kBET in circle.

**Table S1.** The average performance of seven methods based on these four metrics

| Model | Weighted Average |
| --- | --- |
| RAW | 0.61 |
| PRECAST | 0.81 |
| DeepST | 0.67 |
| STAligner | 0.67 |
| GraphST | 0.75 |
| SPIRAL | 0.74 |
| STitch3D | 0.73 |
| Spatialign | 0.50 |
